# Sulfopin, a selective covalent inhibitor of Pin1, blocks Myc-driven tumor initiation and growth *in vivo*

**DOI:** 10.1101/2020.03.20.998443

**Authors:** Christian Dubiella, Benika J. Pinch, Daniel Zaidman, Theresa D. Manz, Evon Poon, Shuning He, Efrat Resnick, Ellen M. Langer, Colin J. Daniel, Hyuk-Soo Seo, Ying Chen, Scott B. Ficarro, Yann Jamin, Xiaolan Lian, Shin Kibe, Shingo Kozono, Kazuhiro Koikawa, Zainab M. Doctor, Behnam Nabet, Christopher M. Browne, Annan Yang, Liat Stoler-Barak, Richa B. Shah, Nick E. Vangos, Ezekiel A. Geffken, Roni Oren, Samuel Sidi, Ziv Shulman, Chu Wang, Jarrod A. Marto, Sirano Dhe-Paganon, Thomas Look, Xiao Zhen Zhou, Kun Ping Lu, Rosalie C. Sears, Louis Chesler, Nathanael S. Gray, Nir London

## Abstract

The peptidyl-prolyl cis-trans isomerase, Pin1, acts as a unified signaling hub that is exploited in cancer to activate oncogenes and inactivate tumor suppressors, in particular through up-regulation of c-Myc target genes. However, despite considerable efforts, Pin1 has remained an elusive drug target. Here, we screened an electrophilic fragment library to discover covalent inhibitors targeting Pin1’s active site nucleophile - Cys113, leading to the development of Sulfopin, a double-digit nanomolar Pin1 inhibitor. Sulfopin is highly selective for Pin1, as validated by two independent chemoproteomics methods, achieves potent cellular and *in vivo* target engagement, and phenocopies genetic knockout of Pin1. Although Pin1 inhibition had a modest effect on viability in cancer cell cultures, Sulfopin induced downregulation of c-Myc target genes and reduced tumor initiation and tumor progression in murine and zebrafish models of MYCN-driven neuroblastoma. Our results suggest that Sulfopin is a suitable chemical probe for assessing Pin1-dependent pharmacology in cells and *in vivo*. Moreover, these studies indicate that Pin1 should be further investigated as a potential cancer target.

## Introduction

Cancer relies on multiple signaling pathways to sustain proliferation and downregulate apoptotic signals ^1^, including the phosphorylation of serine/threonine - proline motif (pSer/Thr-Pro) ^2,3^ found in many cellular proteins. This motif is specifically recognized and isomerized by the peptidyl-prolyl isomerase NIMA-interacting-1 (Pin1), which is the only known phosphorylation-dependent isomerase amongst the ∼30 peptidyl-prolyl cis-trans isomerases (PPIases) in the human proteome ^4^. Pin1-mediated isomerization was shown to impact substrate stability ^5–9^, activity ^10,11^, subcellular localization ^8^, and binding to interaction partners, including Proline-directed kinases and phosphatases, which are mostly *trans*-specific ^12–14^. Thus, Pin1 represents a unified signaling hub that is exploited by cancer to activate oncogenes and inactivate tumor suppressors ^15,16^.

Several lines of evidence suggest that aberrant Pin1 activation drives oncogenesis. Pin1 is over-expressed and/or -activated in at least 38 tumor types ^17^. While elevated Pin1 expression correlates with poor clinical prognosis ^18,19^, polymorphisms that lower Pin1 expression are associated with reduced cancer risk ^20^. Pin1 sustains proliferative signaling in cancer cells by upregulating over 50 oncogenes or growth-promoting factors ^15,16^, including NF-κB ^9^, Notch1 ^21^ and c-Myc ^22^, while suppressing over 20 tumor suppressors or growth-inhibiting factors, such as FOXOs ^23^, Bcl2 ^24^ and RARα ^25^. Furthermore, Pin1-null mice are resistant to tumorigenesis induced by mutant p53 ^26^, activated HER2/RAS ^27^, or constitutively expressed c-Myc ^28^. In addition, Pin1 inhibition sensitizes cancer cells to chemotherapeutics ^25,29,30^, radiation therapy ^31^ and blocks the tumorigenesis of cancer stem cells ^16,32,33^. However, as evidenced by the Cancer Dependency Map ^34^, Pin1 is not essential to cellular viability and Pin1-null mice are viable, though they develop premature aging phenotypes ^7,35^.

Taken together, these studies suggest that pharmacological inhibition of Pin1 has the potential to block multiple cancer-driving pathways simultaneously ^16^ and with limited toxicity. Indeed, compounds that inhibit Pin1, such as juglone ^36^, all-*trans* retinoic acid (ATRA)^37^, arsenic trioxide (ATO)^38^ and KPT-6566 ^39^, exhibit anti-cancer activity and have been used to investigate the role of Pin1 in oncogenesis. Nevertheless, these compounds have been shown to lack specificity and/or cell permeability thus making them unreliable as tools to interrogate the consequences of pharmacological inhibition of Pin1 *in vivo* ^*40–42*^. We have recently developed a selective Pin1 covalent peptide inhibitor ^43^. However, it too, was unsuitable for *in vivo* applications.

The active-site of Pin1 contains a nucleophilic cysteine residue (Cys113) which is suitable for the development of targeted covalent inhibitors ^43–45^. Such covalent inhibitors have several advantages over non-covalent inhibitors ^46–48^ and have been extremely successful against both traditional ^49–51^ and challenging targets ^52–55^.

To explore this strategy, we undertook a covalent Fragment-Based Drug Discovery (FBDD) screening campaign targeting Cys113 in Pin1’s PPIase active site. FBDD, which focuses on low molecular weight compounds (typically < 300 Da), is a successful hit discovery approach ^56–58^ that has led to several drugs and chemical probes ^57,59^. It offers good coverage of chemical space and a high probability of binding due to lower molecular complexity ^60,61^. Covalent FBDD, which increasingly comes to the fore ^62–72^, combines the advantages of FBDD with the improved potency conferred by covalent bond formation.

Optimization of screening hits from this campaign, led to the development of Sulfopin, a double-digit nanomolar, highly selective Pin1 inhibitor that engages Pin1 in cells and *in vivo*. We found that Pin1 inhibition induced modest viability effects in 2D cancer cell culture only after prolonged exposure, and resulted in the downregulation of Myc-dependent target genes. In MYCN-driven models of neuroblastoma, both in zebrafish and in mice, Sulfopin significantly reduced tumor initiation as well as tumor progression. Sulfopin is therefore the first selective Pin1 inhibitor suitable for the evaluation of Pin1 biology *in vivo*, and provides evidence that Pin1 warrants further exploration as a potential anti-cancer target.

## Results

### A covalent fragment screen identifies Pin1 binders

We previously compiled a library of 993 electrophilic fragments featuring mildly reactive ‘warheads’ that can react covalently with cysteines in target proteins ^62^. We screened our library against Pin1 to identify fragment leads for covalent inhibitor development. Purified catalytic domain of Pin1 was incubated with the fragment library (2 μM protein, 200 µM compound; 24 h at 4 °C), followed by intact protein liquid chromatography/mass-spectrometry (LC/MS) to identify and quantify Pin1 labeling (Fig. 1A). In total, 111 fragments irreversibly labeled Pin1 by > 50% under the assay conditions (Fig. 1B; Supp. Dataset 1). Among the 48 top hits (labeling > 75%) nine chloroacetamides shared a common cyclic sulfone core, suggesting a structure-activity relationship (SAR; Fig. 1B, see Supp. Fig. 1 & Supp. Table 1 for a full list of hits containing this motif). Given that the identified sulfone-containing hits were non-promiscuous in previous fragment screens against a diverse panel of proteins ^62^, we selected them for further development. To avoid undesired reactivity arising from the presence of an additional Michael acceptor in the 2-sulfolene fragments, we focused exclusively on the sulfolane analogs.

**Figure 1.**
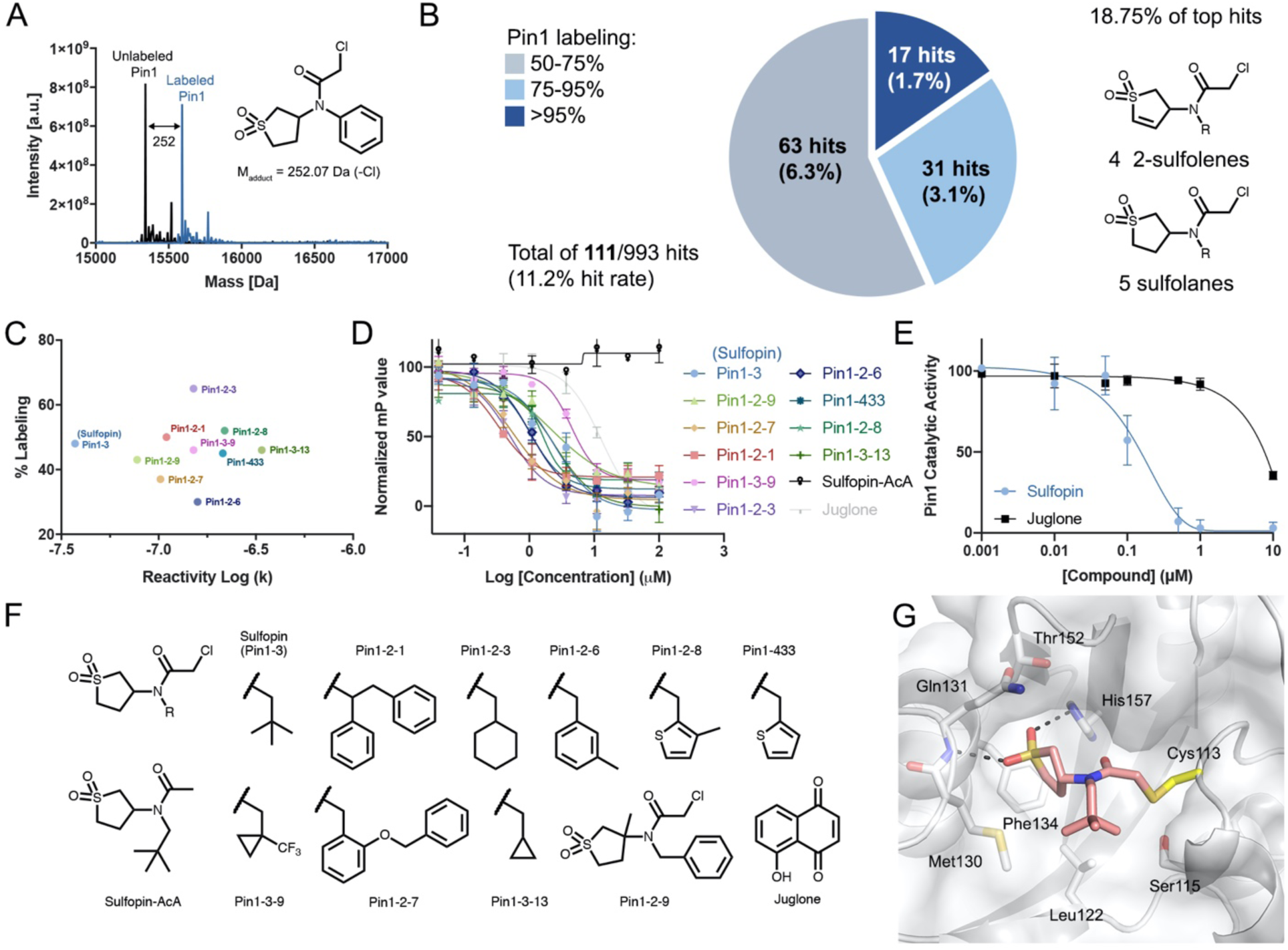
The discovery of a covalent Pin1 binding fragment. **A**. Intact protein LC/MS spectra of Pin1 (black) directly identifies covalent binders (blue) in the electrophilic library screen (200 µM compound for 24 h) **B**. Distribution of hits in the Pin1 screening campaign and their corresponding labeling in percent: Nine hits (18.75%) out of 48 top hits that labeled Pin1 > 75% (dark and light blue) share sulfolene or sulfolane moieties. **C**. 2D-analysis of the top ten optimized binders (structures in F.): Labeling % in the LC/MS-assay is plotted against reactivity (log (k)) suggests Sulfopin for further biological evaluation. **D**. Fluorescence polarization assay with Pin1 and the top ten binders including juglone and a non-reactive control (Sulfopin-AcA), after 14 h pre-incubation. See Supp. Table 3 for apparent K_i_ **E**. Substrate activity assay of Pin1 with Sulfopin and juglone. **F**. Structures of the top ten binders in the Pin1 labeling LC/MS-assay, the non-reactive control Sulfopin-AcA and juglone. **G**. X-ray crystal structure of Pin1 in complex with Sulfopin (1.4 Å resolution, PDB code 6VAJ). Pin1 (white) with relevant side-chains in stick representation. Sulfopin is shown in pink. Hydrogen-bonds are depicted as dashed lines.

### Fragment optimization yields potent Pin1 binding that is not driven by high warhead reactivity

To guide compound optimization, we used the covalent docking program, DOCKovalent ^73^. Docking the sulfolane hits into various Pin1 structures and inspecting highly ranked poses, suggested two plausible binding modes, in which the sulfolane or a lipophilic moiety (‘R’ in Fig. 1B) either protruded into the hydrophobic proline-binding pocket, or interacted with a hydrophobic patch adjacent to Cys113 (Supp. Fig. 2). Both poses suggested maintaining the sulfolane, while diversifying the lipophilic moiety.

To optimize these original hits, we synthesized or purchased a total of 26 sulfolane-containing compounds, featuring a range of aliphatic, arylic, biphenylic or heterocyclic side-chains (Supp. Fig. 3). To identify high-affinity binders, we assessed their irreversible labeling of Pin1 in a second screen under more stringent conditions, using a 1:1 ratio of protein:compound, and a shorter incubation time (2 µM compound; 1 h at RT). Remarkably, 25 out of 26 of these second-generation compounds showed better labeling than the original screening hits, which showed no labeling under these new conditions (Supp. Table 2). Overall, the hits from this second screen revealed that a wide range of lipophilic moieties were tolerated. We concluded that an additional methylene group between the amide and the lipophilic side chain was crucial for Pin1 labeling, as exemplified by four matched molecular pairs that lacked this group and showed no labeling (Supp. Fig. 4). The top ten binders from the second screen (Fig. 1C,F) showed 35-65% Pin1 labeling. We next evaluated these analogs in a competitive fluorescence polarization (FP) assay using a FITC-labeled substrate mimetic peptide inhibitor ^74^. Following a 14 h incubation with recombinant full-length Pin1, all analogs competed in the FP to a greater extent than juglone, an often cited Pin1 inhibitor (Fig. 1D).

Identifying and excluding overly reactive and potentially promiscuous compounds is critical in the development of covalent probes. Accordingly, we assessed the thiol reactivity of the top binders using a high-throughput assay that we previously applied to the entire fragment library ^62^. We found that there was no correlation between labeling efficiency and reactivity (Fig. 1C; Pearson R = 0.003; Supp. Fig. 5). This was particularly evident when comparing Pin1-3, which has a tert-butyl side chain, with the structurally similar Pin-1-13, which has a cyclopropyl side chain. Both compounds showed similar Pin1 labeling (48% and 46%), but their reactivity varied by an order of magnitude, with Pin1-3 being dramatically less reactive. Furthermore, the compound with the highest degree of Pin1 labeling, Pin1-2-3, showed only median reactivity relative to the entire panel.

We have previously shown ^62^ that electrophiles with reactivity rate constants higher than 10^−7^M^-1^s^-1^ may exhibit non-selective cytotoxicity. We evaluated selected compounds from the top Pin1 labelers in a viability assay against IMR90 lung fibroblasts, and Pin1-3 was the only compound that did not show any toxicity up to 25 μM (Supp. Table 3). Pin1-3 has the lowest inherent reactivity of the top identified Pin1 labelers, and does not exhibit non-selective cytotoxicity, therefore showing the best balance of potency and selectivity. For this reason, we selected Pin1-3, henceforth **Sulfopin**, for further evaluation.

### Sulfopin potently binds and inhibits Pin1

Sulfopin displayed potent Pin1 binding in the FP assay ^37^ with an apparent K_i_ = 17 nM (after 14 h; Fig. 2B). A corresponding reversible compound (Sulfopin-AcA; Fig. 1F), which lacks the chloride leaving group, was inactive in the FP assay, suggesting that the binding affinity of Sulfopin is dependent on its electrophile (Fig. 1D,2B). Sulfopin-AcA was subsequently used as a negative control compound. We then performed the FP assay in a dose- and time-dependent manner to determine the K_inact_ of Sulfopin to be 0.03 min^-1^, with a second order rate constant (K_inact_/K_i_) of 84 M^-1^s^-1^ (Supp. Fig. 6). Sulfopin also inhibited the catalytic activity of Pin1 with an apparent K_i_ of 211 nM measured at 12 h, as determined using a chymotrypsin-coupled peptidyl-prolyl isomerization assay ^75^ (PPIase assay; Fig. 1E).

**Figure 2.**
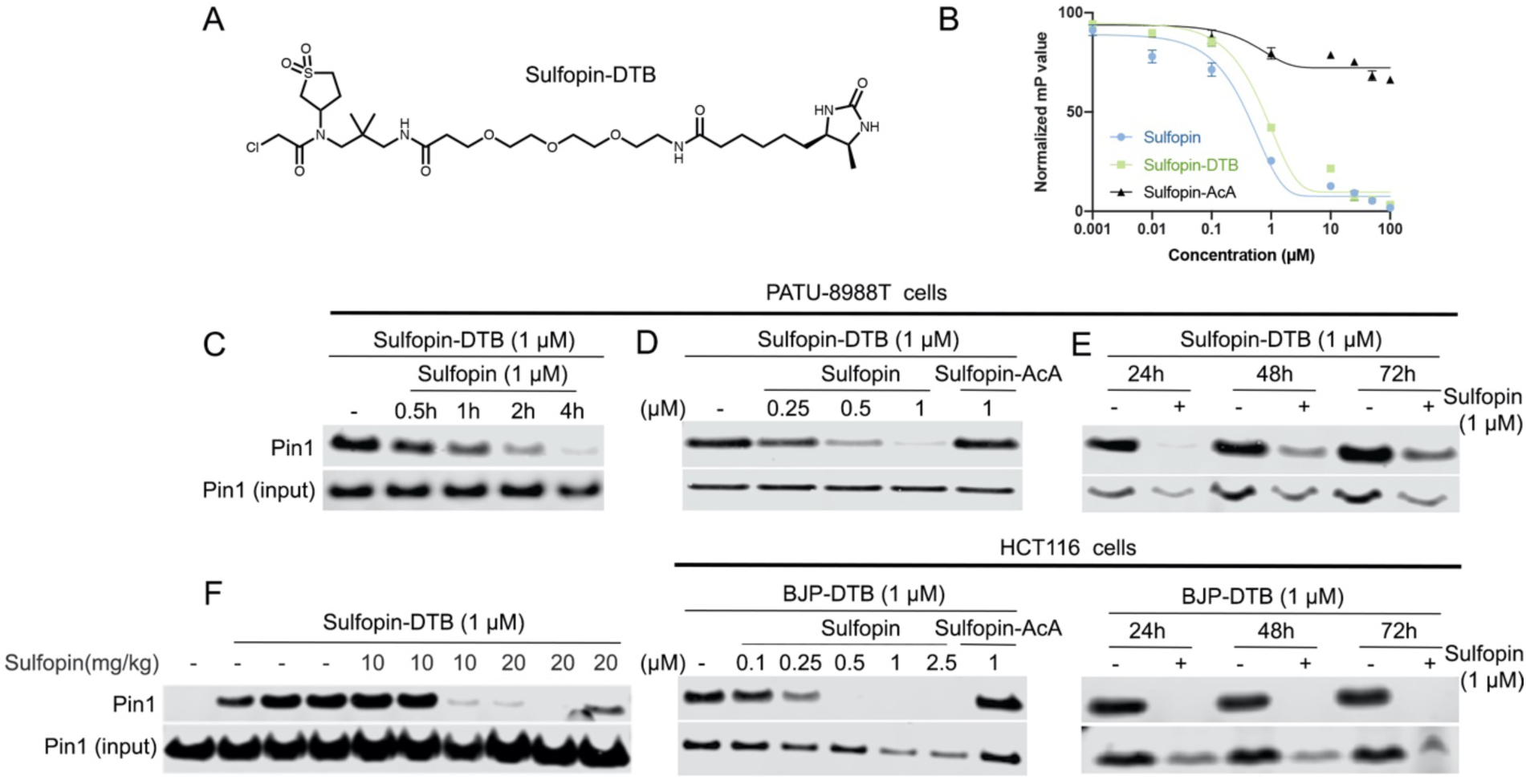
Sulfopin engages Pin1 in cells and *in vivo*. **A**. Chemical structure of the desthiobiotin probe Sulfopin-DTB. **B**. Competitive FP assay shows that Sulfopin-DTB binds Pin1 with similar potency to Sulfopin. **C**. Sulfopin shows time dependent engagement in PATU-8988T cells. Cells were incubated for the indicated times with Sulfopin (1 μM), then lysed and incubated with Sulfopin-DTB (1 μM). Following enrichment, Pin1 levels were analysed by Western blot. **D**. Sulfopin fully engages Pin1 in PATU-8988T cells at 1 μM and in HCT116 cells at 0.5 μM (See Supp. Fig. 9B for a structure of BJP-DTB). Cells were incubated with Sulfopin at the indicated concentration for 5 h, followed by lysis and DTB probe incubation (1 h, 1 μM). The non-covalent control Sulfopin-AcA is not able to engage Pin1. **E**. Sulfopin shows long-term engagement with Pin1. PATU-8988T and HCT116 cells were incubated with or without Sulfopin for the indicated times, followed by lysis and incubation with DTB probes. Significant engagement (> 50-100%) is still evident after 72 h. **F**. Sulfopin engages Pin1 *in vivo*. Mice were treated by oral gavage with the indicated amounts of Pin1, over two days for a total of three doses. Following this treatment, their spleens were lysed for a competition pull-down experiment with Sulfopin-DTB.

To evaluate the binding mode of Sulfopin, we determined the co-crystal structure of Pin1 in complex with Sulfopin at 1.4 Å resolution (Supp. Table 4). The complex structure shows clear electron density to Cys113 in the 2F_O_-F_C_ omit map, which confirmed a covalent interaction (Supp. Fig. 7). In this structure, the sulfolane ring occupies the hydrophobic proline-binding pocket that is formed by Met130, Gln131, Phe134, Thr152 and His157 (Fig. 1G). Furthermore, the sulfonyl oxygens mediate hydrogen bonds to the backbone amide of Gln131, and the imidazole NH of His157. These hydrogen bonds are analogous to those formed between Pin1 and arsenic trioxide ^38^ (Supp. Fig. 8). The tert-butyl group of Sulfopin covers a shallow hydrophobic patch formed by Ser114, Leu122 and Met130, but is mostly solvent-exposed, which explains the broad range of hydrophobic moieties that were tolerated at this position during the optimization campaign.

Despite being a very small ligand (heavy atom count: 17, cLogP: 0.36), Sulfopin efficiently exploits interactions with the active site of Pin1, even in the absence of a negatively charged moiety to interact with the phosphate-binding pocket, thus overcoming the cell permeability issues of previous Pin1 inhibitors, which are often highly anionic ^74,76–78^.

### Sulfopin engages Pin1 in cells and in vivo

To evaluate the target engagement of Sulfopin in cells, we developed a desthiobiotin (DTB)-labeled probe for competition pull-down experiments. Based on the co-crystal structure of Sulfopin bound to Pin1, we derivatized the mostly solvent-exposed tert-butyl group of Sulfopin with a PEG-linked DTB (Sulfopin-DTB; Fig. 2A). Sulfopin-DTB showed similar potency (apparent K_i_ = 38 nM in FP assay; Fig. 2B), and successfully engaged Pin1 in PATU-8988T cell lysates, achieving robust pull-down at 1 μM following a 1 h incubation period (Supp. Fig. 9A).

To assess the cellular target engagement and cell permeability of Sulfopin, we performed a live cell competition assay in PATU-8988T and HCT116 cells. A 1 μM treatment of Sulfopin achieved complete Pin1 engagement within 4 h (with about 50% after 2 h; Fig. 2C), and maintained significant engagement up to 72 h (Fig. 2E). Sulfopin exhibited dose-dependent competition for Pin1 binding, with maximal competition evident at 0.5-1 μM, while the negative control Sulfopin-AcA showed no competition (Fig. 2D). We found that this cellular engagement was extensible to other cell lines, such as IMR32, and MDA-MB-231 (Supp. Fig. 9C,D).

Ultimately, we were interested whether Sulfopin could be used *in vivo*. Sulfopin had encouraging metabolic stability in mouse hepatic microsomes (T_1/2_ = 41 min), prompting us to submit it for pharmacokinetic (PK) profiling. In three mice, oral administration of 10 mg/kg Sulfopin achieved an average C_max_ of 11.5 μM and oral bioavailability (F%) of 30% (Supp. Table 5), suggesting that Sulfopin is suitable for oral *in vivo* dosing. We next evaluated the toxicity of Sulfopin in an acute toxicity model in which mice were administered daily doses of 10, 20 or 40 mg/kg Sulfopin by intraperitoneal (IP) injection for two weeks. No adverse effects or weight loss were recorded, and a post-mortem examination found no readily detectable pathologies.

Using Sulfopin-DTB, we were also able to assess the *in vivo* engagement of Sulfopin. To do this, mice were treated with three doses (over the course of two days) of vehicle, 10, or 20 mg/kg Sulfopin by oral gavage, followed by lysis of the spleens for a competition pull-down experiment. Effective Pin1 engagement was observed in 1 out of 3 mice treated with 10 mg/kg Sulfopin, and in 3 out of 3 mice treated with 20 mg/kg Sulfopin, with target engagement monitored by loss of Sulfopin-DTB-mediated pull-down (Fig. 2F). Based on these results, we chose a 40 mg/kg dose for further mouse experiments to ensure complete Pin1 engagement.

### Sulfopin is highly selective

To evaluate the selectivity of Sulfopin in cells, we assessed its target profile using Covalent Inhibitor Target-site Identification (CITe-Id ^79^, Fig. 3A). This chemoproteomic platform enables the identification and quantification of the dose-dependent binding of covalent inhibitors to cysteine residues on a proteome-wide scale. In this competition experiment, live PATU-8988T cells were incubated with Sulfopin (100, 500, 1000 nM) for 5 h, followed by cell lysis and co-incubation with Sulfopin-DTB (2 μM) for 18 h. Following trypsin digestion and avidin enrichment, the DTB-modified peptides were analyzed by multidimensional LC-MS/MS. Out of 162 cysteine residues labeled by Sulfopin-DTB, only Pin1 Cys113 (Fig. 3B; >2 s.d. from the median; Supp. Dataset 2A) exhibited dose-dependent competition (Fig. 3C), indicating the pronounced selectivity of Sulfopin.

**Figure 3.**
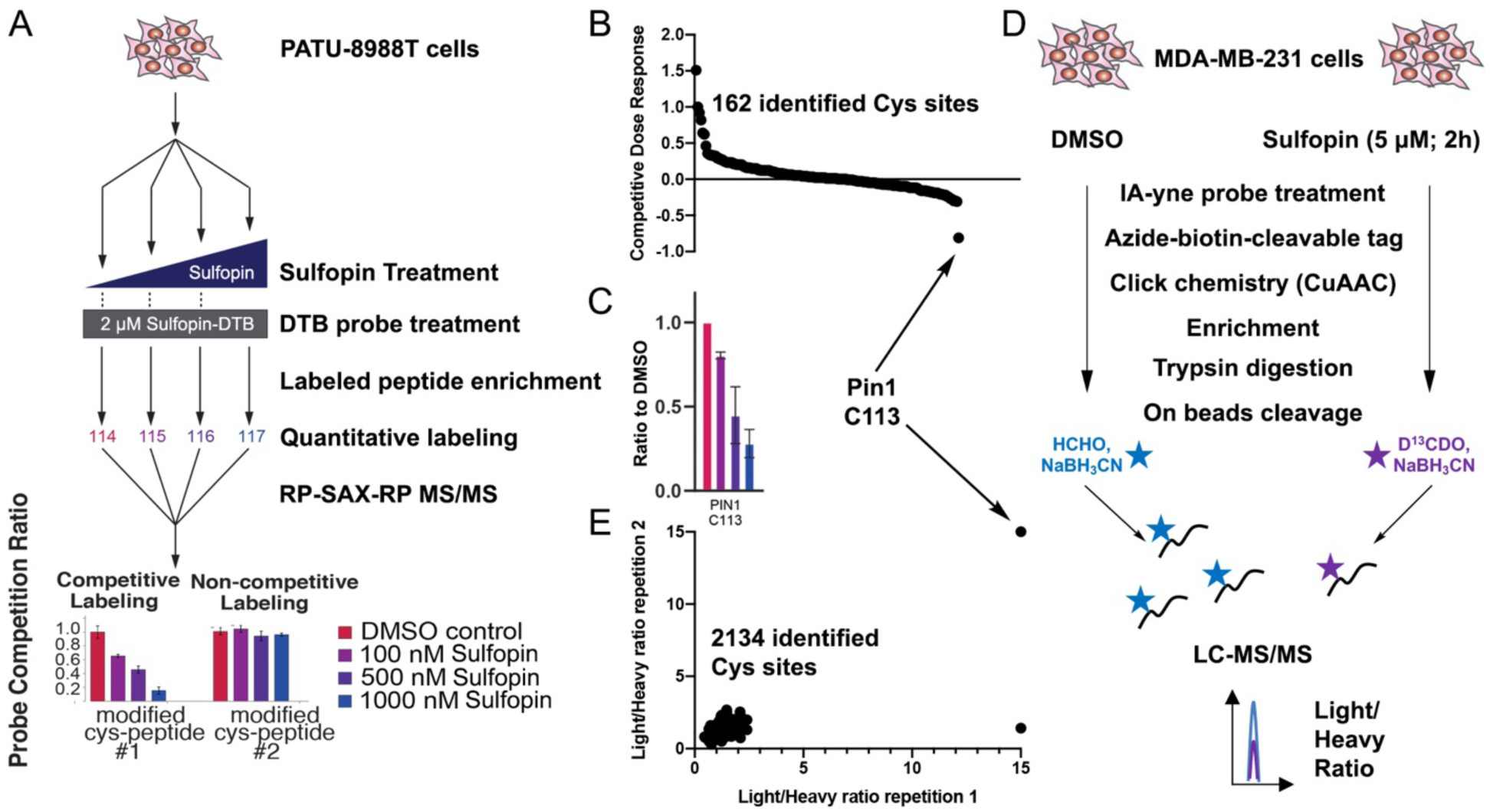
Sulfopin is highly selective for Pin1 in cells. **A**. Scheme for Covalent Inhibitor Target-site Identification (CITe-Id) profiling of competitively labeled cysteine residues across the proteome following dose response treatment with Sulfopin. **B**. CITe-Id results. Out of 162 identified labeled cysteine residues, only C113 in Pin1 is labeled in a dose-dependent manner (see Supp. Dataset 2A for full list of identified peptides). **C**. Dose dependency of C113 labeling in CITe-Id experiment. **D**. Scheme for independent rdTOP-ABPP experiment for assessing Sulfopin proteomic selectivity **E**. Out of 2134 identified cysteines in the experiment, only two cysteines showed light/heavy ratio > 2.5, of these, one cysteine did not replicate, and only Pin1 C113 showed the maximal ratio of 15 in both replicates.

To further investigate Sulfopin’s selectivity, we used a complementary chemoproteomic method. We performed an rdTOP-ABPP experiment to profile its cysteine targets throughout the proteome (Fig. 3D)^80^. This variant of the isoTOP-ABPP technique enables the site-specific quantification of cysteine binding by label-free covalent inhibitors. In brief, MDA-MB-231 cells were treated with Sulfopin (5 μM, 2 h), lysed and labeled with a bioorthogonal Iodoacetamide-alkyne (IA-yne) probe that was then conjugated to a cleavable biotin tag by copper-catalyzed azide−alkyne cycloaddition (CuAAC). After enrichment on beads, the peptides were isotopically derivatized by duplex reductive dimethylation, cleaved and analyzed via LC-MS/MS analysis. We identified Cys113 of Pin1 as the top ranked cysteine that was labeled by Sulfopin at biologically relevant concentration (5 µM), with a competition ratio R = 15 across two biological replicates (Fig. 3E; Supp. Dataset 2B). All other identified cysteines showed R values < 2.5. Overall, we used two independent chemoproteomic techniques in two different cell lines to demonstrate that Sulfopin has exquisite selectivity for Pin1 Cys113 and is therefore a suitable probe with which to interrogate Pin1’s function in cells and in vivo.

### Treatment with Sulfopin phenocopies known Pin1 knockout phenotypes

We next assessed whether pharmacological inhibition of Pin1 by Sulfopin could recapitulate two previously reported phenotypes associated with Pin1 genetic knockout. First, Pin1 knockout was reported to abrogate phosphorylation of IRAK1 Thr209 and resensitize radioresistant cancer cells to irradiation ^31^. Accordingly, we found that treating radioresistant HeLa cells with Sulfopin significantly resensitized them to irradiation in a dose dependent manner (Fig. 4A), and also decreased IRAK1 Thr209 phosphorylation at concentrations as low as 100nM (Fig. 4B,C).

**Figure 4.**
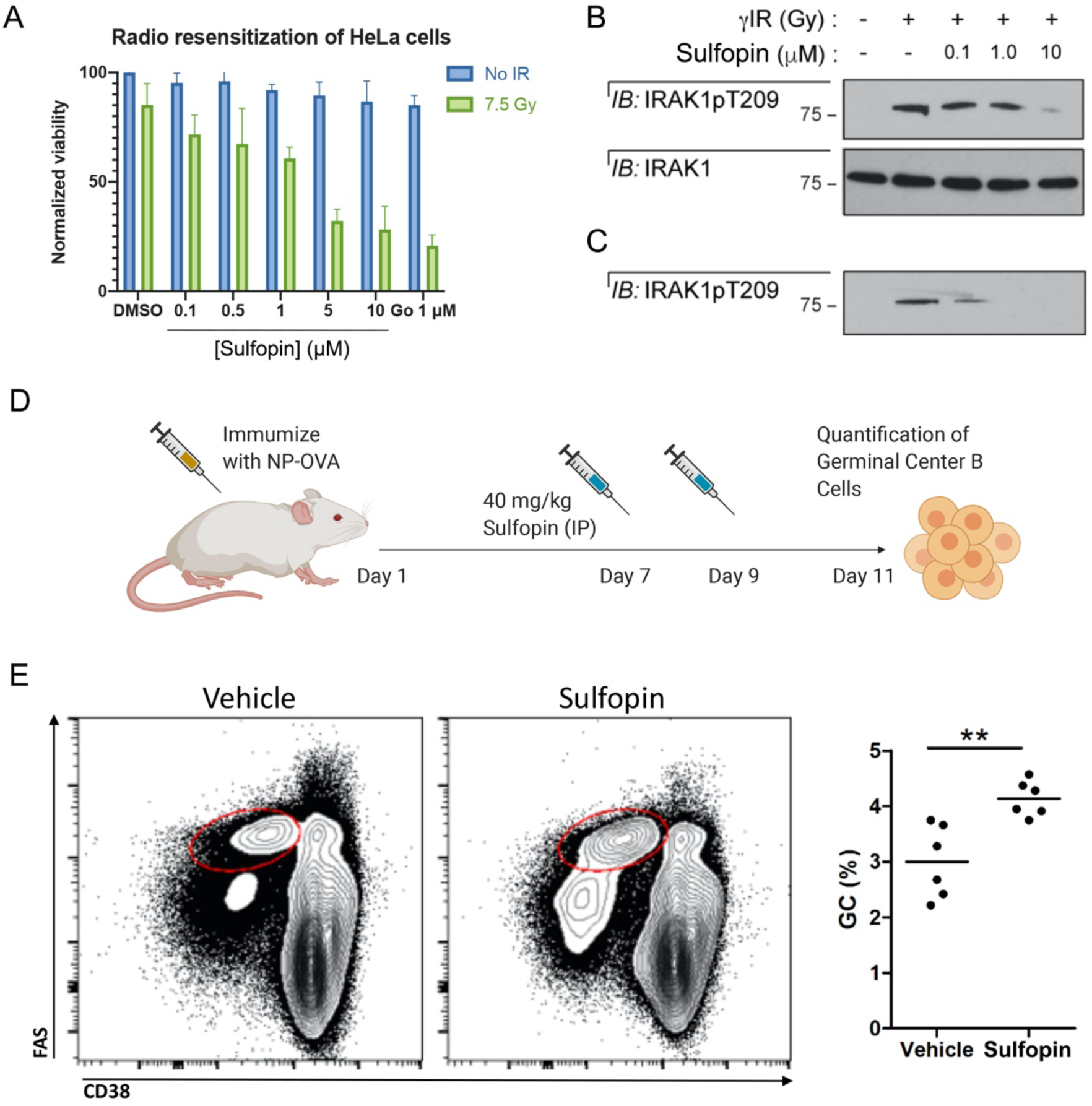
Sulfopin phenocopies Pin1 knockout phenotypes. **A**. HeLa cells were treated with either DMSO, Sulfopin, or Go6976 (a Chk1 inhibitor) and exposed to 7.5 Gy IR 1 h after drug treatment. Viability was assessed 3 days post-IR. Sulfopin shows a dose dependent sensitization of the cells to irradiation. **B**. Western blot analysis was performed 24 h post-IR, showing Sulfopin blocked phosphorylation of Thr209 of IRAK1. **C**. A shorter exposure shows that Sulfopin inhibits IRAK1 phosphorylation already at concentrations of 0.1 μM. **D**. A scheme for testing the effect of Sulfopin *in vivo* on germinal center B cells in response to immunization. **E**. Representative flow cytometric plots with Vehicle and Sulfopin (left) and quantification (right) of FAS^Hi^ CD38^-^ germinal center (GC) cells in WT mice 11 days after immunization with NP-OVA. ** p<0.01, two tailed Student’s t test.

Second, germinal centers (GCs) are sites where B cells proliferate and undergo somatic hypermutation in a BCL6- and Myc-dependent manner. Pin1-deficient mice are reported to display a significant increase in the frequency of GC B cells in response to immunization ^81^. To induce GCs and examine Sulfopin’s effect on GC B cells, we immunized the hind foot pads of 12 wild-type (WT) mice with ovalbumin coupled to the 4-hydroxy-3-nitrophenylacetyl (NP-OVA). The mice were injected with two doses of Sulfopin (IP; 40mg/kg) or vehicle on days 7 and 9 post immunization, at the peak of the GC response, and on day 11 the mice were sacrificed and frequency of GC B cells in lymph nodes was assessed by flow cytometry (Fig. 4D). In accordance with the previous report ^81^, Sulfopin treated mice exhibited 1.34-fold higher proportion of GC B cells compared to mice treated with control vehicle (Fig. 4E). Taken together, these data demonstrate that Sulfopin phenocopies the effects of Pin1 genetic deletion.

### Viability effects of Sulfopin in cancer cell lines

To broadly profile the anti-proliferative activity of Sulfopin, we used the PRISM platform ^82^ (Broad Institute) to evaluate its potency against 300 human cancer cell lines. The PRISM method enables high-throughput, pooled screening of mixtures of cell lines, which are each labeled with a 24-nucleotide barcode ^82^. In all 300 cancer cell lines profiled, Sulfopin demonstrated limited or no anti-proliferative activity after a 5-day treatment, with IC_50_ values > 3 µM. This result aligns with our initial cytotoxicity screening, as well as data from the Cancer Dependency Map, in which Pin1 was not identified as a significant genetic dependency in CRISPR-Cas9 and RNAi screens across hundreds of adherent and suspension cancer cell lines (https://depmap.org/portal/). This suggests that the strong single-agent cytotoxicity of previously published Pin1 inhibitors, such as juglone, is likely attributable to off-target effects ^42,83^ (Supp Fig. 10).

We next assessed whether Sulfopin treatment might induce antiproliferative activity effects after prolonged treatments (6-8 days). To ensure that target engagement was maintained for the duration of the experiment, we replenished Sulfopin in fresh media every 48 h. We found that Sulfopin treatment impacted the viability of PATU-8988T cells in a Pin1-dependent manner, having no effect on proliferation in the corresponding Pin1 KO cells (Fig. 5A). To evaluate whether this time-dependent growth phenotype was extensible to other cancer types, we performed additional experiments in breast (MDA-MB-468), prostate (PC3), and ovarian (Kuramochi) cancer cell lines. MDA-MB-468 cells showed the most pronounced sensitivity to Sulfopin, which diminished cell viability in a dose- and time-dependent manner, while Sulfopin-AcA did not affect cell proliferation (Fig. 5B). Both PC3 and Kuramochi cells exhibited only very modest sensitivity to Sulfopin treatment.

**Figure 5.**
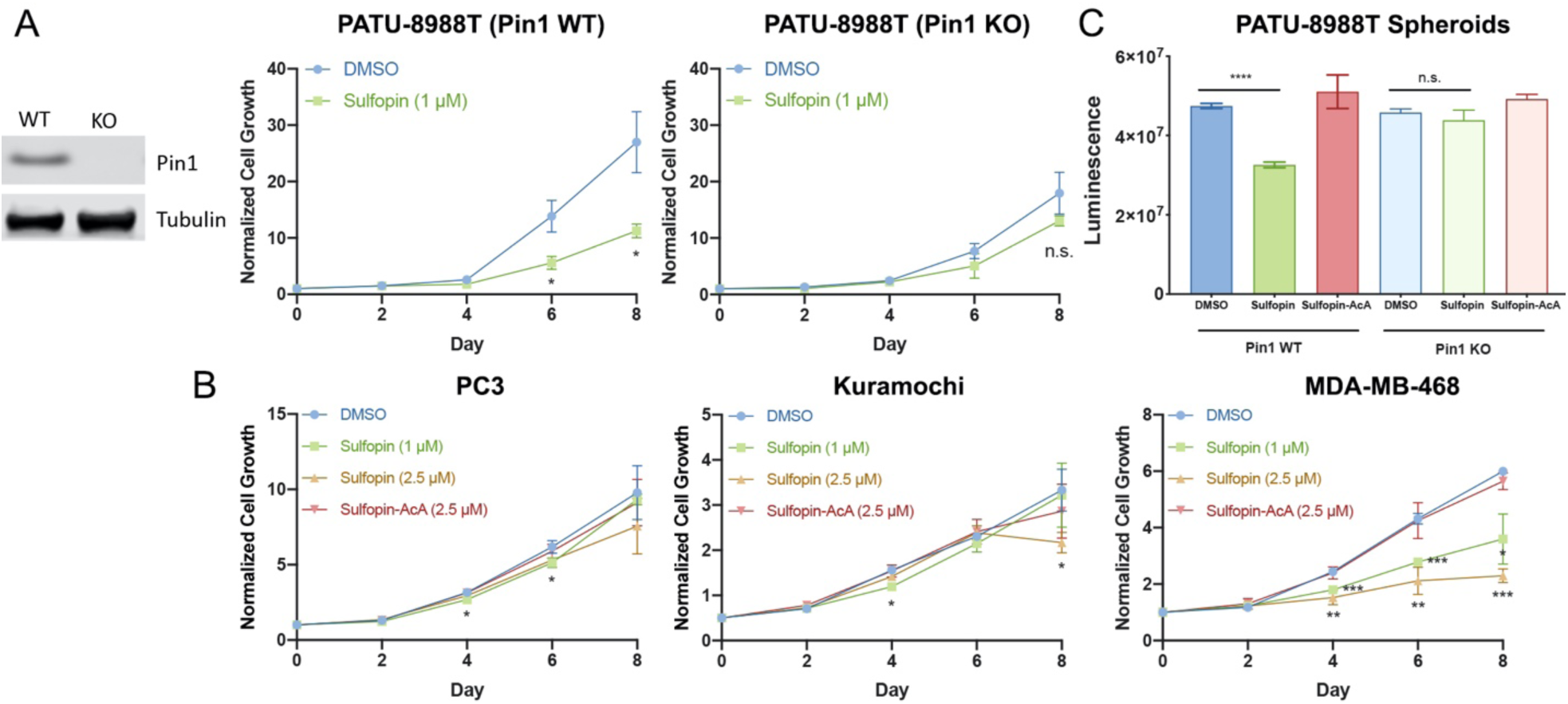
Sulfopin has a Pin1 dependent viability effect on long term exposure. **A**. We generated a PATU-8988T Pin1 knockout (KO) cell line, as evidenced by Western blot. Sulfopin (1 μM) has a significant effect on cellular viability after 6 and 8 days in the WT PATU-8988T cells, but shows no significant effect on viability in the Pin1 KO cells (n=3). **B**. Sulfopin shows varying anti-proliferative effects across cancer cell lines, with the most pronounced sensitivity observed in MDA-MB-468 cells (n=3). **C**. Relative viability of PATU-8988T WT and Pin1 KO cells grown in 100% Matrigel domes following treatment with Sulfopin (1 μM; n=9) or Sulfopin-AcA (1 μM; n=9). The non-covalent control Sulfopin-AcA shows no effect in any of the tested systems. Statistical significance calculated using Student’s t test with unequal variance (n.s. = p > 0.05; * = p < 0.05; ** = p < 0.01; *** = p < 0.001; **** = p < 0.0001). In all graphs, error bars indicate standard deviation.

Given that three-dimensional (3D) culture models can reflect *in vivo* results better than monolayer cell culture^84^, we next evaluated the anti-proliferative activity of Sulfopin in PATU-8988T WT or Pin1 KO cells grown in 3D matrigel domes. Following a 9-day treatment, replenishing compound in media every 3 days, Sulfopin demonstrated modest anti-proliferative activity in PATU-8988T WT Pin1 cells, with no effects observed in PATU-8988T Pin1 KO cells, suggesting an on-target phenotype (Fig. 5C).

Collectively, these data suggest that Pin1 inhibition does not cause proliferation defects at short time points, but that it does moderately affect cell proliferation after prolonged treatments, and in 3D cell culture.

### Sulfopin downregulates Myc transcriptional activity

We and others have previously shown that Pin1 regulates the c-Myc oncoprotein ^85^, affecting Myc protein stability ^86^ as well as its DNA binding and transcriptional activity ^22,87^. Pin1 physically interacts with c-Myc ^86,88^, altering isomerization of P63 following phosphorylation of S62. We have shown that overexpression of Pin1 can lead to an increase in the transcription of c-Myc target genes while knockdown of Pin1 decreases Myc-dependent transcription ^22^.

To test whether Sulfopin affects Myc transcriptional output, we treated Mino B cells with Sulfopin (1 μM; 6 h; in triplicates) or DMSO and performed a global RNA sequencing analysis to detect differentially expressed genes as the result of this perturbation. 206 genes were found to be significantly down-regulated (Fig. 6A; Supp. Dataset 3). We performed a gene set enrichment analysis of these genes using Enrichr ^89^, against a dataset of genes identified by ChIP-seq (chromatin immunoprecipitation followed by sequencing) for various transcription factors. Myc target genes in K562 cells and HeLa-S3 cells appeared as the most enriched set and the third most enriched set, respectively (adjusted p-value of 1.99×10^−16^ and 2.00×10^−13^ respectively; Fig. 6B), validating a significant downregulation of Myc’s transcriptional signature by Sulfopin. To further validate the effect of Sulfopin on Myc transcriptional activity, we co-transfected HEK-293 cells with a Myc reporter construct (4x-Ebox-Luciferase) and Pin1. As expected, Pin1 expression increased Myc transcriptional activity, while treatment with 2 μM Sulfopin for 48 h resulted in a significant reduction in relative luciferase activity as compared to DMSO control (Fig. 6C). Together these results suggest that treatment with Sulfopin downregulates Myc target genes and, thus, points to Myc-driven cancers as natural candidates for its therapeutic application. Accordingly, we next evaluated Sulfopin in *in vivo* models of MYCN-driven neuroblastoma (NB).

**Figure 6.**
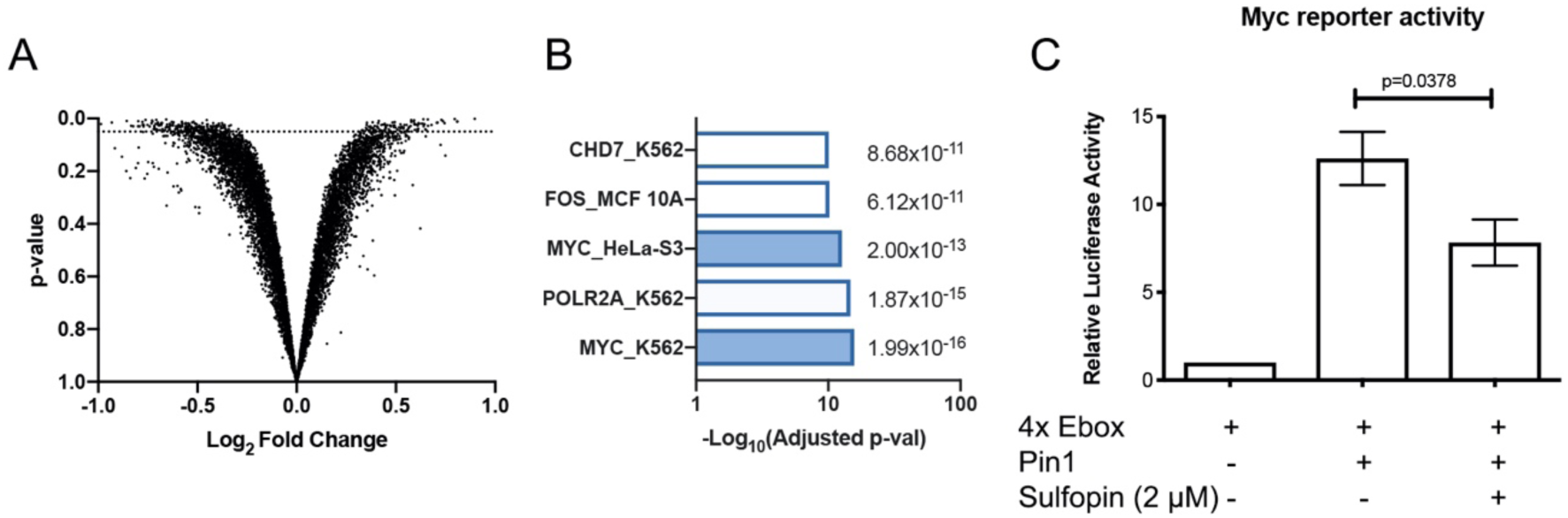
Sulfopin downregulates Myc transcription. **A**. Results of an RNA-sequencing experiment, comparing the change in RNA levels between Mino B cells treated with either Sulfopin (1 μM, 6 h, in triplicates) or DMSO. Each dot presents the Log_2_ fold change of a transcript (x-axis) vs. the p-value for significance of that change (Student’s t test; y-axis). The dotted line indicates p=0.05. 206 genes were significantly down regulated. **B**. Results of a gene set enrichment analysis using Enrichr against the ENCODE TF ChIP-seq set. Two of the most enriched sets are Myc target genes from different cell lines. **C**. HEK293 cells were transfected with a 4x Ebox-luciferase reporter for Myc transcriptional activity levels. Co-transfection with Pin1 increased reporter activity, while 48 h treatment with Sulfopin significantly (Student’s t test) reduced this activity compared to DMSO.

### Sulfopin blocks tumor initiation and progression in MYCN-driven zebrafish models of neuroblastoma

To evaluate Sulfopin’s effects on Myc-driven cancers, we turned to a zebrafish model of neuroblastoma ^90–92^, a pediatric malignancy derived from the peripheral sympathetic nervous system (PSNS)^93^. During the development of normal zebrafish embryos, neural crest-derived PSNS neuroblasts form the primordial superior cervical ganglia (SCG) and intrarenal gland (IRG) at the age of 3 to 7 days post fertilization (dpf), can be visualized using the dβh:EGFP fluorescent reporter ^91^ (Fig. 7A). Overexpression of the MYCN oncogene, which is the oncogenic driver in approximately 20% of human high-risk neuroblastomas, causes the fish to develop neuroblast hyperplasia in the PSNS of Tg(dβh:MYCN;dβh:EGFP) transgenic zebrafish (Fig. 7A upper right). These neuroblast hyperplasia rapidly progress into fully transformed tumors that faithfully resemble human high-risk neuroblastoma ^90–92^. When Sulfopin was added to the fish water containing Tg(dβh:MYCN;dβh:EGFP) zebrafish (at concentrations of 25-100 µM), the neuroblastoma-initiating hyperplasia was significantly suppressed and fully transformed neuroblastoma did not develop over the treatment period (Fig. 7A,B). This indicates that Sulfopin blocks neuroblastoma initiation *in vivo* in this tumor model. In addition, no evidence of toxicity was observed in embryos treated with Sulfopin at these concentrations, further supporting our findings in mice that Sulfopin is well tolerated by healthy tissues *in vivo*.

**Figure 7.**
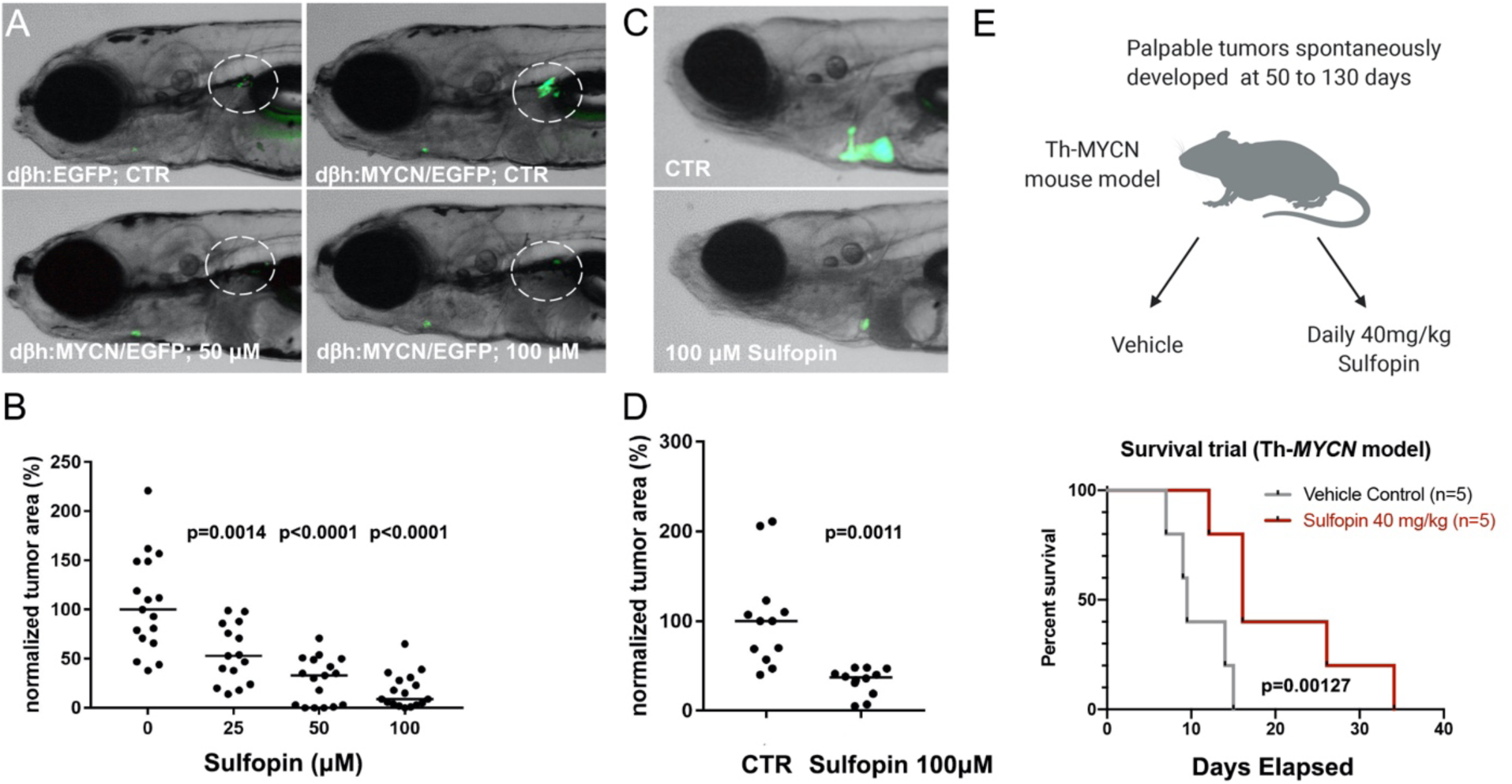
Sulfopin abrogate MYCN driven neuroblastoma growth *in vivo*. **A**. PSNS cells in the primordial superior cervical ganglia (SCG) and intrarenal gland (IRG; highlighted by dotted circles) in representative embryos of Tg(dβh:EGFP) (upper left) and Tg(dβh:MYCN;dβh:EGFP) (upper right) transgenic zebrafish at 7 dpf. Representative 7 dpf Tg(dβh:MYCN;dβh:EGFP) zebrafish treated with 50 µM Sulfopin (bottom left) and 100 µM Sulfopin (bottom right). The primordial SCG plus IRG areas are highlighted by dotted circles. **B**. Quantification of GFP+ cells in the primordial SCG and IRG of 7 dpf Tg(dβh:MYCN;dβh:EGFP) embryos treated with Sulfopin at multiple doses. A Mann-Whitney test with confidence intervals of 95% was used for the analysis of significance (p-value) and the quantitative data are reported as median. **C**. Representative zebrafish embryos transplanted with neuroblastoma cells isolated from four-month old Tg(dβh:MYCN;dβh:EGFP) donor zebrafish and treated with DMSO control (upper) or 100 µM Sulfopin added to the fish water (lower). **D**. Quantification of EGFP-positive tumor area in zebrafish embryos treated with DMSO control and 100 µM Sulfopin added to the fish water. A Mann-Whitney test with confidence intervals of 95% was used for the analysis of significance (p-value) and the quantitative data are reported as median. **E**. Survival trials in Th-*MYCN* mice. Mice were randomly assigned to treatment groups, and treated daily with either vehicle or 40 mg/kg Sulfopin, as soon as the tumors were palpable. Treated mice showed significant (p=0.0127) increase in overall survival, with an average increase of 10 days survival.

We then assessed the anti-tumor activity of Sulfopin on the maintenance of fully transformed neuroblastoma cells *in vivo* in primary tumor derived allograft (PDA) models constructed in transplanted zebrafish embryos. EGFP-labeled neuroblastoma cells were dissected from four-month-old Tg(dβh:MYCN;dβh:EGFP) donor zebrafish, disaggregated, counted and 200-400 GFP-labeled tumor cells were injected intravenously into the Duct of Cuvier (common cardinal vein) of 2 dpf zebrafish embryos ^94^. One day after injection, 100 µM Sulfopin or the DMSO control was added to the fish water containing embryos bearing the transplanted EGFP-labeled neuroblastoma cells. After five days, we quantified the area of the EGFP-labeled tumor mass in treated embryos, and discovered that tumor masses in the DMSO-treated embryos grew larger over the five days of treatment, while the tumor masses decreased in size in the Sulfopin treated embryos (Fig. 7C,D). Hence, Sulfopin treatment not only suppressed MYCN-driven neuroblastoma initiation, but also suppressed the growth and survival *in vivo* of transplants of fully transformed primary neuroblastoma tumor cells.

### Sulfopin blocks tumor initiation and progression in a MYCN-mouse model of neuroblastoma

Following the encouraging results in the zebrafish models, we also assessed the effects of Sulfopin in a murine model of neuroblastoma - the Th-MYCN GEM (genetically engineered mouse) model of neuroblastoma, in which human MYCN is expressed under the tyrosine hydroxylase promoter ^95^. The Th-MYCN faithfully recapitulate the major molecular and histopathologic features of high-risk MYCN-amplified neuroblastoma, and the model has been widely used for therapeutics studies. The study was performed using both male and female hemizygous mice, which spontaneously developed palpable tumors at 50 to 130 days with a 25% penetrance. Once tumors were palpable, mice were randomly assigned to treatment groups, and treated once per day with either vehicle or 40 mg/kg Sulfopin. In this aggressive model, we assessed the overall survival of the mice. Treated mice showed significant (p=0.0127) increase in overall survival, with an average increase of 10 days for Sulfopin treated mice (Fig. 7E).

## Discussion

Despite extensive, decades-long efforts to discover Pin1 inhibitors, no approach has yielded a compound capable of selectively blocking Pin1 *in vivo*. Here, we describe the development and *in vivo* characterization of Sulfopin, a highly selective and potent, covalent Pin1 inhibitor with low inherent reactivity and negligible toxicity, which blocks tumor initiation and progression in both zebrafish and mouse models of neuroblastoma.

While inhibitors such as juglone, ATRA, ATO and KPT-6566 pioneered the investigation of Pin1 in cancer-related contexts, they all exert their anti-cancer effects in part through Pin1-independent mechanisms ^41^ that include DNA-damage ^39^, induction of Pin1 degradation ^37^ or by directly blocking transcription ^83^. Our recently reported covalent peptide inhibitor BJP-06-005-3 ^43^, while selective in cells, is not suitable for *in vivo* application. In contrast, Sulfopin is highly specific for Pin1, as we established here using multiple orthogonal experimental strategies. Therefore, this highly selective *in vivo* tool allowed us to investigate Pin1 as a *bona fide* drug target in cancer for the first time. Thus far, Pin1 has proven a challenging target for both ligand-^74^ and structure-based ^76–78^ approaches, even in studies employing high-throughput-screenings of up to one million compounds ^76^. This is in part due to the shallow nature of the Pin1 active site, which is evolved to bind peptides, further complicated by the phosphate binding site, which favors negatively charged moieties. To overcome this hurdle, we screened a library of electrophilic fragments, which can compensate for sparse protein interfaces by irreversibly binding to the Pin1 Cys113. The screen resulted in a relatively high hit rate of 11% (111 compounds with >50% labeling) compared to previous screens ^62^. This might be explained by the high reactivity of Cys113, which was found amongst the most reactive cysteines in several chemoproteomic campaigns ^67^.

A major finding during our hit optimization campaign was that intrinsic warhead reactivity of Pin1 binders did not correlate with potency (Supp. Fig. 5). Despite the high structural similarity of the optimized compounds, their reactivities spanned over 30-fold (Supp. Table 2), with Sulfopin having the lowest reactivity of the best binders (Fig. 1C), with reactivity nearing that of acrylamides (Supp. Fig. 11). This low reactivity manifested in exquisite cellular selectivity. Two separate chemoproteomic approaches, performed in two different cell lines, both identified Pin1 as the sole target of Sulfopin by a wide margin (Fig. 3). This result is rare for covalent inhibitors ^79^ and even for FDA approved covalent drugs ^96^.

This proteomic selectivity for Pin1 allowed us to show that short-term Pin1 inhibition does not induce cytotoxicity, despite previous results reported with less selective inhibitors ^39,97–102^. Juglone, for instance, has identical LD_50_ for cell lines lacking Pin1 ^42,43^ (Supp. Fig. 10). Sulfopin induced little to no cytotoxicity across a panel of 300 cancer cell lines (PRISM screen), though it significantly impacted cancer cell growth after prolonged treatments (6-8 days; Fig. 5A,B) and in 3D-cell culture (Fig. 5C). The irreversible binding of Sulfopin allowed us to demonstrate target engagement in cells and *in vivo* (Fig. 2), showing that its favorable PK/PD profile (Supp. Table 5) translated to complete *in vivo* target engagement and target modulation as demonstrated by the ability of Sulfopin to phenocopy previous genetic results *in vivo* (Fig. 4). This result underscores the utility of Sulfopin as an *in vivo* tool compound, with broad applicability to domains such as immunology, and as we later show, cancer biology.

In agreement with previous studies ^22,86^, our RNA sequencing experiment suggested Myc target gene downregulation as a major consequence of Pin1 inhibition (Fig. 6A,B). This conclusion is supported by transcriptional suppression in the Myc luciferase reporter assay (Fig. 6C), and the demonstrated efficacy of Pin1 inhibition in various MYCN-driven cancer models (Fig. 7). However, given that Pin1 plays a central role in numerous signaling pathways, it is likely that Myc-downregulation is not the only mechanism at play. Indeed, additional transcription factors showed significant downregulation in the RNA-seq dataset, including RelA (Supp. Dataset 4) which was also previously linked to Pin1 ^9,103^ and could contribute to oncogenesis. Other pathways that were previously linked to Pin1 have also been implicated in neuroblastoma ^104,105^. RNA-seq experiments in additional cell lines (and perhaps additional time points) might shed more light on other processes regulated by Pin1.

Further investigation is needed to establish whether the therapeutic efficacy of Sulfopin can be further enhanced. For instance, we have yet to reach dose limiting toxicity with Sulfopin, and observed dose-dependent responses in zebrafish (Fig. 7B), indicating that treatment with higher doses might enable more pronounced effects in mice. Furthermore, its favorable toxicity profile could enable the use of Sulfopin in combination with other drugs. Since Pin1 regulates numerous cellular pathways, it stands to reason that Pin1 inhibition may be synthetic lethal with the loss of other targets. Ongoing studies will clarify the full therapeutic potential of Sulfopin.

In summary, we present a potent and selective, covalent Pin1 inhibitor with *in vivo* activity and no toxicity that allows investigation of Pin1 biology in physiologically relevant models in health and disease. Using Sulfopin as a pharmacological tool will enable the study of the many diverse processes that are regulated by Pin1.

## Methods

### Electrophile library screening

993 compounds as 20 mM DMSO stocks in 384 well-plate format were transferred into a 384 well plate working copy by combining 0.5 μL of five compounds per well. The catalytic domain of Pin1 (2 µM) in 20 mM Tris, 75 mM NaCl, pH7.5 was incubated with 200 μM for each compound and moderately shaken for 24 h at 4°C. The reaction was stopped by the addition of formic acid to 0.4% final concentration. The LC/MS runs were performed on Waters ACUITY UPLC class H, in positive ion mode using electrospray ionization. UPLC separation using C4 column (300 Å, 1.7 µM, 21 mm×100 mm). the column was held at 40°C, and the autosampler at 10°C. Mobile solution A was 0.1% formic acid in water and mobile phase B was 0.1% formic acid in acetonitrile. Run flow was 0.4 mL/min. Gradient used for BSA was 20% B for 2 minutes increasing linearly to 60% B for 4 minutes holding at 60% B for 2 minutes, changing to 0% B in 0.1 minutes and holding at 0% for 1.9 minutes. Gradient for the other proteins was 20% B for 2 minutes increasing linearly to 60% B for 3 minutes holding at 60% B for 1.5 minutes, changing to 0% B in 0.1 minutes and holding at 0% for 1.4 minutes. The mass data was collected at a range of 600-1300 m/z. Desolvation temperature was 500°C with a flow rate of 1000 liter/hour. The voltage used were 0.69 kV for the capillary and 46 V for the cone. Raw data was processed using openLYNX and deconvoluted using MaxEnt. Labeling assignment was performed as previously described^62^.

### Covalent Docking

Covalent docking was performed using DOCKovalent 3.7^73^ against 16 structures of Pin1. PDB codes: 1PIN, 2ITK, 2Q5A, 2XP3, 2ZQV, 2ZR4, 3IK8, 3KAB, 3KCE, 3NTP, 3ODK, 3OOB, 3TC5, 3TCZ, 3TDB, 3WH0. The docked compounds include seven sulpholane hits from the electrophilic library with the following IDs: PCM-0102138, PCM-0102178, PCM-0102105, PCM-0102832, PCM-0102313, PCM-0102760, PCM-0102755. The covalent bond length was set to 1.8Å and the two newly formed bond angles to Cβ-Sγ-C=109.5 ± 5° and Sγ-C-Ligatom=109.5 ± 5°.

### Thiol reactivity assay

50 μM DTNB was incubated with 200 μM TCEP in 20 mM sodium phosphate buffer pH 7.4, 150 mM NaCl, for 5 minutes at room temperature, in order to obtain TNB^2-^. 200 μM compounds were subsequently added to the TNB^2-^, followed by immediate UV absorbance measurement at 412 nm at 37°C. The UV absorbances were acquired every 15 minutes for 7 h. The assay was performed in a 384 well-plate using a Tecan Spark10M plate reader. Background absorbance of compounds was subtracted by measuring the absorbance at 412 nm of each compound in the same conditions without DTNB. Compounds were measured in triplicates. The data was fitted to a second order reaction equation such that the rate constant k is the slope of ln([A][B0]/[B][A0]). Where [A0] and [B0] are the initial concentrations of the compound (200 μM) and TNB^2-^ (100 μM) respectively, and [A] and [B] are the remaining concentrations as a function of time as deduced from the spectrometric measurement. Linear regression using Prism was performed to fit the rate against the first 4 h of measurements.

### Pin1 expression and purification

A construct of full-length human Pin1 in a pET28 vector was overexpressed in E. coli BL21 (DE3) in LB medium in the presence of 50 mg/ml of kanamycin. Cells were grown at 37°C to an OD of 0.8, cooled to 17°C, induced with 500 μM isopropyl-1-thio-D-galactopyranoside, incubated overnight at 17°C, collected by centrifugation, and stored at −80°C. Cell pellets were sonicated in buffer A (50 mM hepes 7.5, 500 mM NaCl, 10% glycerol, 20 mM Imidazole, and 7 mM BME) and the resulting lysate was centrifuged at 30,000 xg for 40 min. Ni-NTA beads (Qiagen) were mixed with lysate supernatant for 30 min and washed with buffer A. Beads were transferred to an FPLC-compatible column and the bound protein was washed with 15% buffer B (50 mM hepes 7.5, 500 mM NaCl, 10% glycerol, 250 mM Imidazole, and 3 mM BME) and eluted with 100% buffer B. Thrombin was added to the eluted protein and incubated at 4°C overnight. The sample was concentrated and passed through a Superdex 200 10/300 column (GE helathcare) in a buffer containing 20 mM HEPES, pH 7.5, 150 mM NaCl, 5% glycerol, and 1 mM TCEP. Fractions were pooled, concentrated to approximately 37 mg/ml and frozen at −80°C.

### Pin1 crystallization and soaking

Apo protein at a final concentration of 1 mM was crystallized by sitting-drop (200 nL + 200 nL) vapor diffusion at 20°C in the following crystallization buffer: 3 M NH_4_SO_4_, 100 mM HEPES-pH7.5, 150 mM NaCl, 1% PEG400, and 10 mM DTT. A volume of 200 nL of 1 mM Sulfopin was added directly to crystals for soaking at 20°C for 16 h. Crystals were transferred briefly into crystallization buffer containing 25% glycerol prior to flash-freezing in liquid nitrogen.

### Crystallization data collection and structure determination

Diffraction data from complex crystals were collected at beamline 24ID-C of the NE-CAT at the Advanced Photon Source at the Argonne National Laboratory. Data sets were integrated and scaled using XDS^106^. Structures were solved by molecular replacement using the program Phaser^107^ and the search model PDB entry 1PIN. Iterative manual model building and refinement using Phenix^108^ and Coot^109^ led to the final models (Supp. Table 4).

### Fluorescence polarization (FP) assay

Binding affinity to Pin1 was determined using a fluorescence polarization (FP) assay to assess competition with an N-terminal fluorescein-labeled peptide (peptide core structure: Bth-D-pThr-Pip-Nal), which was synthesized by Proteintech. The indicated concentrations of candidate compounds were pre-incubated for 12 h at 4°C with a solution containing 250 nM glutathione S-transferase (GST)-Pin1, 5 nM of fluorescein-labeled peptide probe, 10 μg/ml bovine serum albumin, 0.01% Tween-20 and 1 mM DTT in a buffer of 10 mM HEPES, 10 mM NaCl and 1% glycerol (pH 7.4). Measurements of FP were made in black 384-well plates (Corning) using an EnVision reader. K_i_ values obtained from the FP assay results were derived from the Kenakin K_i_ equation: Kenakin K_i_ = (Lb)(EC_50_)(K_d_)/(Lo)(Ro) + Lb(Ro–Lo + Lb–K_d_), where K_d_ [M]: K_d_ of the probe, EC_50_ [M]: concentration of unlabeled compound that results in 50% inhibition of binding (obtained from FP assay), total tracer Lo [M]: probe concentration in FP, bound tracer Lb [M]: 85%, fraction of probe bound to Pin1, total receptor Ro [M]: Pin1 concentration in the FP assay, as described^37,110^.

### Pin1 PPIase activity assay

Inhibition of Pin1 isomerase activity was determined using the chymotrypsin-coupled PPIase assay, using GST-Pin1 and Suc-Ala-pSer-Pro-Phe-pNA peptide substrate, as described previously^75^. GST-Pin1 was pre-incubated with the indicated concentrations of compound for 12 h at 4°C in buffer containing 35 mM HEPES (pH 7.8), 0.2 mM DTT, and 0.1 mg/mL BSA. Immediately before the assay was started, chymotrypsin (final concentration of 6 mg/mL) was added, followed by the addition of the peptide substrate (Suc-Ala-pSer-Pro-Phe-pNA peptide substrate, final concentration 50 mM). The *K*_i_ value obtained from the PPIase assay was derived from the Cheng–Prusoff equation, *K*_i_ = IC_50_/ (1 + *S*/K_m_), where K_m_ is the Michaelis constant for the peptide substrate, *S* is the initial concentration of the substrate in the assay, and IC_50_ is the half-minimal inhibitory concentration of the inhibitor.

### Cell Culture and Reagents

PATU-8988T (DSMZ), MDA-MB-468 (ATCC), and HeLa (ATCC) cells were cultured in Dulbecco’s modified Eagle’s medium (DMEM) supplemented with 10% fetal bovine serum (FBS, Sigma) and 1% penicillin/streptomycin. PC3 (ATCC), IMR32 (ATCC) and Kuramochi (Panagiotis A. Konstantinopoulos’s laboratory) cells were cultured in RPMI 1640 medium with L-glutamine, supplemented with 10% FBS and 1% penicillin/streptomycin. All cell lines were cultured at 37 °C in a humidified chamber in the presence of 5% CO_2_. All cell lines were tested for the absence of Mycoplasma infection on a monthly basis.

### Immunoblotting

Whole cell lysates for immunoblotting were prepared by pelleting cells from each cell line at 4°C (300 g) for 5 minutes. The resulting cell pellets were washed 1x with 5 mL ice-cold PBS and then resuspended in the indicated cell lysis buffer. Lysates were clarified at 14,000 rpm for 15 minutes at 4°C prior to quantification by BCA assay (Pierce, cat#23225). Whole cell lysates were loaded into Bolt 4-12% Bis-Tris Gels (Thermo Fisher, cat#NW04120BOX) and separated by electrophoreses at 95 V for 1.5 h. The gels were transferred to a nitrocellulose membrane using the iBlot Gel Transfer at P3 for 6 minutes (Thermo Fisher, cat#IB23001) and then blocked for 1 h at room temperature in Odyssey blocking buffer (LICOR Biosciences, cat#927-50010). Membranes were probed using antibodies against the relevant proteins at 4°C overnight in 20% Odyssey Blocking Buffer in 1x TBST. Membranes were then washed three times with 1x TBST (at least 5 minutes per wash) followed by incubation with the IRDye goat anti-mouse (LICOR, cat#926-32210) or goat anti-rabbit (LICOR, cat #926-32211) secondary antibody (diluted 1:10,000) in 20% Odyssey Blocking Buffer/1x TBST for 1 h at room temperature. After three washes with 1x TBST (at least 5 minutes per wash), the immunoblots were visualized using the ODYSSEY Infrared Imaging System (LICOR). Antibodies used against various proteins were as follows: Pin1 (1:1,000, Cell Signaling cat#3722), α-Tubulin (1:1,000, Cell Signaling cat#3873), IRAK1 (Cell Signaling Technology #4504), IRAK1 pT209 (Assay Biotech #A1074).

### Lysate Pull-Down with Sulfopin-DTB

The indicated cells were lysed in 50 mM Hepes, pH 7.4, 1 mM EDTA, 10% glycerol, 1 mM TCEP, 150 mM NaCl, 1 mM EDTA, 0.5% NP-40, and protease inhibitor tablet (Roche cat#4693159001). After clarifying (14,000 rpm for 15 min), lysates were incubated with the indicated concentrations of Sulfopin-DTB for 1 h at 4°C, using 500 μg of lysate per sample. Lysates were then incubated with streptavidin agarose resin (30 μL of 1:1 beads:lysis buffer slurry) (Thermo scientific, cat#20349) for 1.5 h at 4°C. Beads were washed four times with 500 μL of buffer (50 mM Hepes, pH 7.5, 10 mM NaCl, 1 mM EDTA, 10 % glycerol), then pelleted by centrifugation and dried. The beads were boiled for 5 minutes at 95°C in 30 μL of 2x LDS + 5% β-mercaptoethanol. Lysates were probed for specified proteins by Western blotting using the Bolt system (Life Technologies).

### Cellular Target Engagement – Competition with Sulfopin-DTB

The indicated cell lines were plated in 10 cm plates with 2.5 million cells per plate in 6 mL of media. The day after plating, cells were treated with DMSO or the indicated concentrations of candidate inhibitor for the indicated time points. The cells were then washed two times with cold PBS (1 mL per 10 cm plate) and collected by scraping with a cell scraper. Cells were lysed in 50 mM Hepes, pH 7.4, 1 mM EDTA, 10% glycerol, 1 mM TCEP, 150 mM NaCl, 1 mM EDTA, 0.5% NP-40, and protease inhibitor tablet (Roche) – using 210 μL of cell lysis buffer per 10 cm plate of cells. After clarifying (14,000 rpm for 15 min), 9 μL of each lysate sample was combined with 4x LDS + 10% β-mercaptoethanol (in a ratio of 3:1), boiled for 5 minutes, and set aside for the input loading control (later to be loaded directly on the gel). Then, 200 μL of each lysate sample was incubated with 1 μM of Sulfopin-DTB for 1 h at 4°C and processed as in “lysate pull-down with Sulfopin-DTB” (above).

### Radiosensitization studies

AlamarBlue-based cell viability assays were performed as previously described^31^. Briefly, HeLa cells were seeded at a density of 200 cells/well in a 96-well plate. After 16 h, cells were treated with Sulfopin and exposed to 7.5 Gy IR using an X-RAD 320 PRECISION X-RAY irradiator 1 hour after drug treatment. At 3 days post-IR, cells were incubated with AlamarBlue (ThermoFisher) at a final concentration of 10%. At 4 days post-IR, absorbance was measured at a wavelength of 570 nm with a 600 nm reference wavelength. Relative fluorescence was calculated using cell-free wells as a control reference, and percentage survival was calculated by comparing with DMSO-treated, non-irradiated controls. Sulfopin efficacy was assessed at 24 h post-IR by Western blot using anti-IRAK1 (Cell Signaling Technology #4504) and anti-IRAK1pT209 (Assay Biotech #A1074) antibodies.

### In vivo germinal centers evaluation

WT mice (C57BL/6) were provided by Harlan, Israel. All experiments with mice were approved by the Weizmann Institute IACUC committee. Mice were immunized by injection of OVA coupled to the hapten 4-hydroxy-3-nitrophenylacetyl (NP-OVA) precipitated in alum (Imject® Alum, Thermo Scientific) into the hind footpads (25ul). Single cell suspensions were obtained by forcing popliteal lymph nodes through a 70 μm mesh into ice cold FACS buffer (EDTA 1 mM and 2% serum in PBS). Cells were incubated with 2 µg/ml anti-16/32 (clone 93) for blockage of FC receptors for 5-10 min. Cell suspensions were washed and incubated with fluorescently-labeled antibodies (B220 V500, FAS FITC, CD38 Alexa fluor 700; Biolegend) for 20-40 min. GC cells were gated as live/single, B220^+^ CD38^Lo^ FAS^Hi^. Cell suspensions were analyzed by Cytoflex (Beckman) flow cytometer.

### Cell Viability Assays: Growth Over Time in 2D-Adherent Monolayer Cell Culture

The indicated cell lines were plated at a density of 500 cells per well (except PATU-8988T cells, which were plated at 100 cells per well to avoid over-confluence by day 8) in 100 μL of media in a 96-well white clear bottom plate (Corning cat#3903), with one plate per time point (Day 0, 2, 4, 6, 8). Cells were treated the day after plating with 1 μL of DMSO, Sulfopin, or Sulfopin-AcA to give the indicated concentrations, and were then incubated at 37°C 5% CO_2_. Every 48 h, the media was aspirated and replaced with fresh media containing fresh compound or DMSO. When the indicated time points had been reached, cell viability was evaluated using the CellTiter-Glo Luminescent Cell Viability Assay (Promega cat#G7570) according to the manufacturer’s standards, measuring luminescence using an Envision plate-reader. The Day 0 time point plates were read the day after plating, prior to compound treatment. N=3 biological replicates were used for each treatment condition.

### Cell Viability Assay: 5 day treatment

PATU-8988T cells were plated in flat bottom 96-well plates (Corning cat #3903) at a density of 1,000 cells per well in 100 μL media and were treated the next day with 1 μL of the indicated compounds in a three-fold dilution series. The cells were incubated at 37°C 5% CO_2_ for 5 days. Anti-proliferative effects were assessed by CellTiter-Glo Luminescent Cell Viability Assay (Promega cat #G7570) according to the manufacturer’s standards, measuring luminescence using an Envision plate-reader. N=3 biological replicates were used for each treatment condition.

### Assessing Antiproliferative Activity in PATU-8988T 3D Cell Culture

Prepare PATU-8988T (WT or Pin1^-/-^) Matrigel suspensions by resuspending cells in 100% cold Matrigel (kept on ice). Plate 1 dome per well in 24-well plate (Greiner CELLSTAR), with 50 μL of cells/Matrigel suspension per dome, and 1,000 cells per dome. After plating the domes, place 24-well plate on top of T175 (previously filled with autoclaved water) in the incubator, let solidify for 15 min. Keeping the 24-well plate on the T175 filled with water, move to the TC hood and carefully add 500 μL of cold DMEM (+10% FBS/1% P/S) per well, then place 24-well plate in incubator. The next day, treat with DMSO, Sulfopin (1 μM), or Sulfopin-AcA (1 μM). Every 3 days, carefully aspirate off the media, add 500 μL of fresh media and retreat. After 9 days, aspirate off the media, add 300 μL of 3D CellTiter-Glo (Promega cat#G9681) per well, shake plate for 1 h. Measure luminescence using an Envision plate-reader. N=3 biological replicates were used for each treatment condition.

### RNA sequencing

Mino cells (acquired from the ATCC) were grown at 37°C in a 5% CO_2_ humidified incubator and cultured in RPMI-1640 (Biological industries), supplemented with 15% Fetal bovine serum (biological industries) and 1% pen-strep solution (biological industries). 11×10^6^ cells were incubated with 1 μM Sulfopin (0.02% DMSO) or with 0.02% DMSO in triplicates for 6 h. Total RNA was isolated with RNeasy kit (Qiagen). RNA libraries were prepared from 2 μg total RNA using SENSE mRNA-Seq library prep kit V2 (lexogen). Total RNA and library quality was analyzed using Qubit fluorometric and TapeStation analysis (Agilent). Samples were sequenced using NextSeq 500/550 High Output Kit v2.5 (illumina) on NextSeq550.

RNA-seq reads were aligned to the human genome (hg19 assembly) using STAR^111^ and gene expression was determined using RSEM^112^ and RefSeq annotations. Differential expression was computed using DESeq2^113^ with default parameters. Genes with baseMean >50 that were downregulated with P<0.05 were further analysed using Enrichr^89^.

### Profiling of Sulfopin-DTB reactive cysteines by CITe-Id

PATU-8988T cells were cultured in DMEM supplemented with 10% Fetal Calf Serum. DMSO (Control) or Sulfopin were diluted in fresh media (final DMSO concentration < 0.1%, final Sulfopin concentration: 100 nM, 500 nM and 1 µM) and added to sub-confluent cultures. After 5 h of incubation at 37°C, cells were harvested using a cell scraper and centrifuged at 300 × g for 3 minutes at 4°C. Cell pellets were washed with ice-cold PBS and centrifuged again. A total of 3 washes were performed before freezing cell pellets at −80°C. This procedure was performed twice on cells independently cultured a week apart. Frozen pellets were then processed essentially as described^79^, except that pre-cleared lysates were treated with 2 µM Sulfopin-DTB overnight at 4 °C. Protein desalting, digestion, enrichment of desthiobiotin modified peptides, iTRAQ stable isotope labeling, peptide clean-up, and multidimensional LC/MS-MS analysis were then performed exactly as described^79^. Data processing and database search was performed as described^79^, except that spectral processing accounted for Sulfopin-DTB specific fragment ions, and Sulfopin-DTB labeling of cysteine was considered as a variable modification by Mascot (version 2.6.2). Inhibitor concentrations and ratios were used to generate a trendline for each labeled site with the slope corresponding to the competitive dose response for each modified cysteine site.

### Profiling of Sulfopin reactive cysteines by rdTOP-ABPP

MDA-MB-231 cells were cultured at 37°C under 5% CO_2_ atmosphere in DMEM culture medium supplemented with 10% FBS and 1% PS. Cells were grown to 70% confluence and incubated with DMSO or 5 μM Sulfopin for 2 h with serum-free medium. Cells were harvested, lysed by sonication in ice-cold PBS containing 0.1% TritonX-100 and centrifuged at 100,000g for 30 min to remove cell debris. Then protein concentrations were determined by BCA protein assay. Proteomes were normalized to 2 mg/mL in 1 mL for each sample.

Each of the DMSO and Sulfopin incubated proteomes was treated with 100 μM iodoacetamide-alkyne (IAyne) for 1 h at room temperature. The proteomes were then reacted with 1 mM CuSO4, 100 μM TBTA ligand, 100 μM biotin-acid-N3 tag and 1 mM TCEP for 1 h. After click reaction, the proteomes were centrifuged at 8000g for 5 min, then the precipitated proteins were washed for two times using cold methanol. The proteomes were resuspended in 1.2% SDS/PBS and diluted to 0.2% SDS/PBS. Finally, the samples were prepared, analyzed on LC-MS/MS and quantified according to the published rdTOP-ABPP protocol^80^. Briefly, the beads from trypsin digestion were washed and resuspended in 100 μL of TEAB buffer. 8 μL of 4% D^13^CDO or HCHO was added to the Sulfopin or DMSO sample respectively. At the same time, 8 μL of 0.6 M NaBH_3_CN was added and the reaction was lasted for 2 h at room temperature. Then the beads were washed again and the modified peptides were cleaved by 2% formic acid. LC-MS/MS data was analyzed by ProLuCID^114^ with static modification of cysteine (+57.0215 Da) and variable oxidation of methionine (+15.9949 Da). The isotopic modifications (+28.0313 and +34.0631 Da for light and heavy labeling respectively) are set as static modifications on the N-terminal of a peptide and lysines. Variable modification on cysteines is set at +322.23688 Da. The ratios were quantified by the CIMAGE software^115^.

### Myc luciferase reporter assay

HEK-293 cells were maintained in DMEM supplemented with 10% standard fetal bovine serum (FBS), 2.5 mM L-glutamine, NEEA and 1x penicillin/streptomycin. Cells were passaged to 80% confluence in 6-well plates and transfected with 4x-Ebox-Luc and Pin1-Flag plasmids as indicated and β-galactosidase as an internal control^22^ using Lipofectamine 3000 (Invitrogen, Carlsbad, CA) following manufacturer’s protocol. Sulfopin treatments were performed at the time of transfection and cells were harvested 48 h later.

Cell were washed with PBS, and then lysed in 1X cell lysis buffer (Promega, Madison, WI). Lysates were sonicated for 10 pulses at an output = 1 and 10% duty (Branson), and incubated on ice for 20 min. Lysates were then cleared by centrifugation at 14K rpm for 10 min at 4°C. Luciferase activity was measured using the Promega Luciferase Assay Kit and Berthold luminometer (Bundoora, Australia) and normalized to β-galactosidase activity ^116^.

### Zebrafish Neuroblastoma Models

All zebrafish studies and maintenance of the animals were performed in accordance with Dana-Farber Cancer Institute IACUC-approved protocol #02-107. For *in vivo* drug treatment, 3-day-old zebrafish embryos were placed in 48-well plates with 5 embryos per well, and treated with DMSO control or Sulfopin in standard egg water.

For neuroblastoma transplantation, zebrafish neuroblastoma cells were harvested by dicing 4-month-old Tg(dβh:MYCN;dβh:EGFP) transgenic zebrafish in PBS. The cell suspension was filtered with Falcon 40-µm cell strainer (Corning, Corning, NY, USA) and loaded into thin-wall borosilicate glass capillary needles (1.0 mm OD, 0.75 mm ID; World Precision Instruments, Sarasota, FL, USA). The recipients, 2-day-old zebrafish Casper embryos, were manually dechorionized and anaesthetized with 0.003% tricaine (Sigma) before being positioned on a 10 cm Petri dish coated with 1% agarose. Intravenous tumor transplantation was performed as described^94^, with 200∼400 cells injected into the Duct of Cuvier of each recipient. One day later, the 3-day-old recipients were randomly divided into 48-well plates and treated with DMSO control or Sulfopin in standard egg water for five days, with drug refreshment on the second day of the treatment.

A Nikon SMZ1500 microscope equipped with a Nikon digital sight DS-U1 camera was used for capturing both the bright field and fluorescent images from live zebrafish and embryos. For PSNS and neuroblastoma quantification, all animals in the same experiments were imaged under the same conditions and the acquired fluorescent images were quantified using ImageJ software by measuring the area of the EGFP fluorescent tumor mass.

### Mice studies (PK/PD/Tox)

PK data was obtained as fee-for-service from Scripps Florida. For the toxicity study 18 mice were used for the following arms: 3 control mice injected with vehicle, 3 mice 10 mg/kg Sulfopin injected once daily, 3 mice 20 mg/kg Sulfopin injected once daily, 3 mice 20 mg/kg Sulfopin injected once every other day, 3 mice 40 mg/kg Sulfopin injected once daily, 3 mice 40 mg/kg Sulfopin injected once every other day. Formulation was (5% NMP, 5% solutol, 20% DMSO). Mice were treated for 14 days.

PD Study was performed at the Dana-Farber Cancer Institute, and the procedure was approved by IACUC under protocol 16-015. Mice were treated for 3 total doses spanning 2 consecutive days with vehicle, or Sulfopin (10 or 20 mpk) by oral gavage, after which the organs were harvested 4 h after the last dose. The spleen of each mouse was ground and lysed in 300 μL of 50 mM Hepes, pH 7.4, 1 mM EDTA, 10% glycerol, 1 mM TCEP, 150 mM NaCl, 1 mM EDTA, 0.5% NP-40, and protease inhibitor tablet (Roche). After clarifying (14,000 rpm for 15 min), the lysates were normalized by BCA and diluted to a final concentration of 2.5 μg/μL. 200 μL of 2.5 μg/μL of each spleen sample was then incubated with 1 μM of Sulfopin-DTB for 1 h at 4°C and processed as in “lysate pull-down with Sulfopin-DTB” (above). To prepare the input samples, each original lysate sample (prior to pull-down) was combined with 4x LDS + 10% β-mercaptoethanol (in a ratio of 3:1), boiled for 5 minutes, and 25 μg of each sample was then loaded on the gel.

### Murine Neuroblastoma models

All experiments were approved by The Institute of Cancer Research Animal Welfare and Ethical Review Body and performed in accordance with the UK Home Office Animals (Scientific Procedures) Act 1986, the United Kingdom National Cancer Research Institute guidelines for the welfare of animals in cancer research^117^ and the ARRIVE (animal research: reporting *in vivo* experiments) guidelines^118^.

Transgenic Th*-MYCN* mice were genotyped to detect the presence of the human *MYCN* transgene^119^. The study was performed using both male and female hemizygous mice, which developed palpable tumors at 50 to 130 days with a 25% penetrance. Tumor development was monitored weekly by palpation by an experienced animal technician. Mice with palpable tumors at >/= 4-5 mm were then enrolled into two groups. Group 1: Animals will all received Sulfopin at 40mg/kg, once per day by oral gavage. Group 2: Animals will all received Vehicle (5% NMP, 5% kolliphor, 20% DMSO) once per day by oral gavage. Studies were terminated when the mean diameter of the tumor reached 15 mm. Tumor volumes were measured by Vernier caliper across two perpendicular diameters, and volumes were calculated according to the formula: *V* = 4/3π [(d1 + d2) / 4]^3^. Mice were housed in specific pathogen-free rooms in autoclaved, aseptic microisolator cages (maximum of four mice per cage).

## Acknowledgments

N.L. is the incumbent of the Alan and Laraine Fischer Career Development Chair; N.L. would like to acknowledge funding from the Israel Science Foundation (grant No. 2462/19), The Rising Tide Foundation, The Israel Cancer Research Fund, the Israeli Ministry of Science. Technology (grant No. 3-14763), and the Moross integrated cancer center. N.L. is also supported by the Helen and Martin Kimmel Center for Molecular Design, Joel and Mady Dukler Fund for Cancer Research, the Estate of Emile Mimran and Virgin JustGiving and the George Schwartzman Fund. C.D. was supported by the Minerva Fellowship program of the Max Planck Society, funded by the German Federal Ministry for Education and Research. The work was supported in part by NIH grant R01CA205153 to K.P.L., N. S. G. and X. Z. Z. N.S.G. was also supported by the Hale Center for Pancreatic Research. Y.C. & C.W. thank the Computing Platform of the Center for Life Science at Peking University for supporting the proteomic data analysis. S.D.P acknowledges funding from the Linde Family Foundation. Part of this research was conducted at the Advanced Photon Source on the Northeastern Collaborative Access Team beamlines (NIGMS P41 GM103403) and SBGrid compiled software (eLife 2013;2:e01456). J.A.M. acknowledges support from the National Institutes of Health (CA233800) and the Mark Foundation for Cancer Research. B.J.P. was supported by the Ruth L. Kirschstein NRSA Individual Predoctoral Fellowship (F31 CA225066), the Training Grant in Chemical Biology (NIH 5 T32 GM007306-41), the Training Grant in Pharmacological Sciences (NIH 5 T32 GM095450-04, B.J.P.), and the Chleck Foundation (B.J.P./Z.M.D.). B.N. was supported by an American Cancer Society Postdoctoral Fellowship (PF-17-010-01-CDD) and the Katherine L. and Steven C. Pinard Research Fund (N.S.G. and B.N.). L. C. is supported by the Cancer Research UK Program Grant (C34648/A18339 and C34648/A14610). Y. J. is supported by a Children with Cancer UK Research Fellowship (2014/176). R.C.S. acknowledges funding support from NCI R01s CA196228 and CA186241 and foundation support from The Brenden-Colson Center for Pancreatic Care. We thank Igor Ulitski for help with RNA-seq analysis and Milka Kostic for critical reading of the manuscript.

