## Supplementary Figures, Tables, and Methods for "Sulfopin, a selective covalent inhibitor of Pin1, blocks Myc-driven tumor initiation and growth *in vivo*"

#### 2-Sulfolenes:

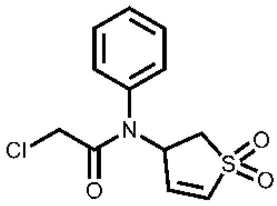

PCM-0102372

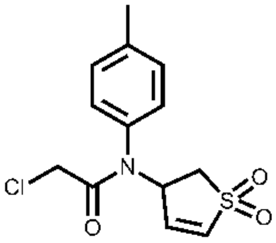

PCM-0102539

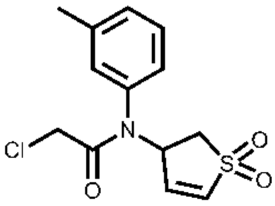

PCM-0102579

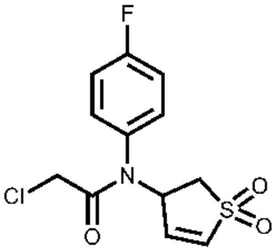

PCM-0102868

#### Sulfolanes:

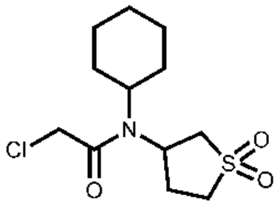

PCM-0102760

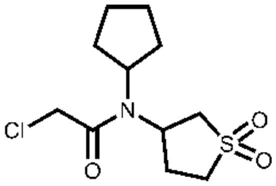

PCM-0102313

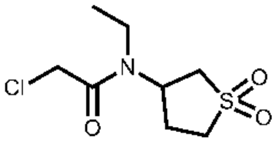

PCM-0102105

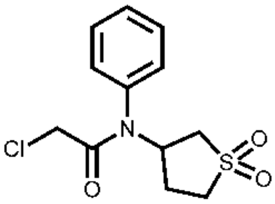

PCM-0102755

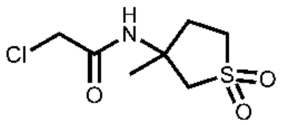

PCM-0102230

**Supplementary Figure 1.** Structures of 2-sulfolenes and sulfolane hits that labeled >75% in the primary screen.

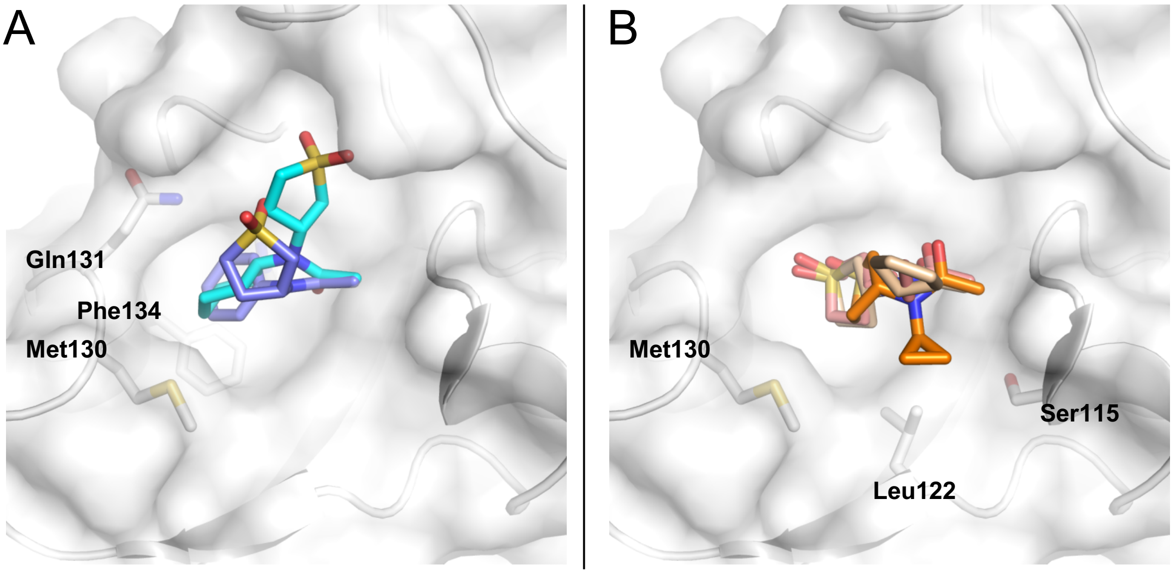


**Supplementary Figure 2. Examples of docking poses with selected screening compounds bound to Pin1** (pdb code: 2ZQV, 2.5 Å).
**A.** The phenyl- and cyclohexyl residues of PCM-010275 (purple) and PCM-010276 (cyan), respectively, protrude into a hydrophobic cavity build up by Met130, Gln131 & Phe134. **B**. The cyclopropyl of PCM-010283 (orange) covers a shallow hydrophobic patch formed by Ser115, Leu122 and Met130. The ethyl- and the cyclopropyl moiety of PCM-010210 (brown) and PCM-0120231 (light brown), respectively, protrude into the solvent.

**Aliphatic:**

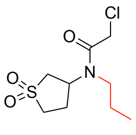
Pin1-028

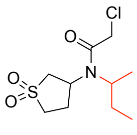
Pin1-053

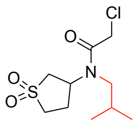
Pin1-3-15

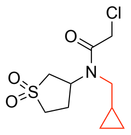
Pin1-3-13

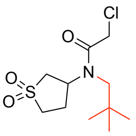
Pin1-3

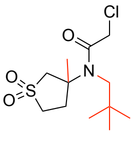
Pin1-18

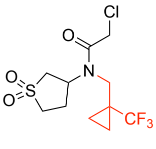
Pin1-3-9

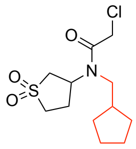
Pin1-3-14

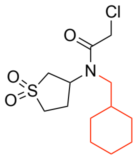
Pin1-2-3

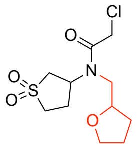
Pin1-838

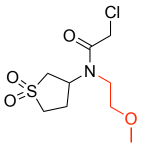
Pin1-324

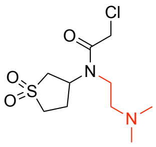
Pin1-707

**Arylic:**

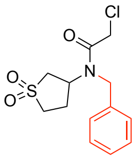
Pin1-437

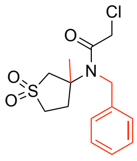
Pin1-2-9

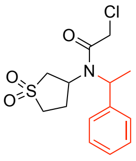
Pin1-2.2

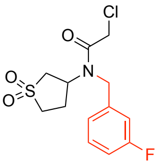
Pin1-2-5

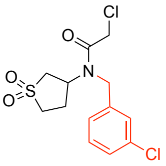
Pin1-2-11

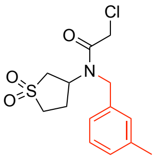
Pin1-2-6

**Bulky arylic/biphenylic**

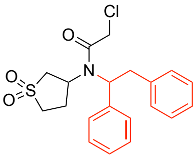
Pin1-2-1

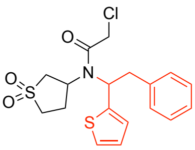
Pin1-2-4

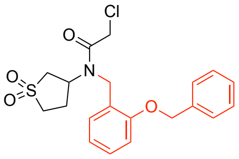
Pin1-2-7

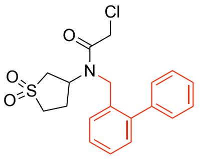
Pin1-2-10

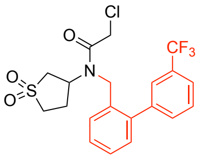
Pin1-3-8

**Heterocyclic:**

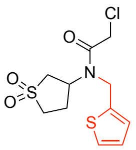
Pin1-433

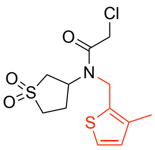
Pin1-2-8

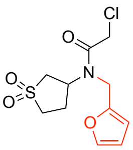
Pin1-128

**Supplementary Figure 3. Sulfolane analogs tested for Pin1 irreversible labeling.** The varying lipophilic residues are denoted in red.

**Sulfolane fragments with no Pin1 labeling at 2 µM (protein:compound) 1h.**

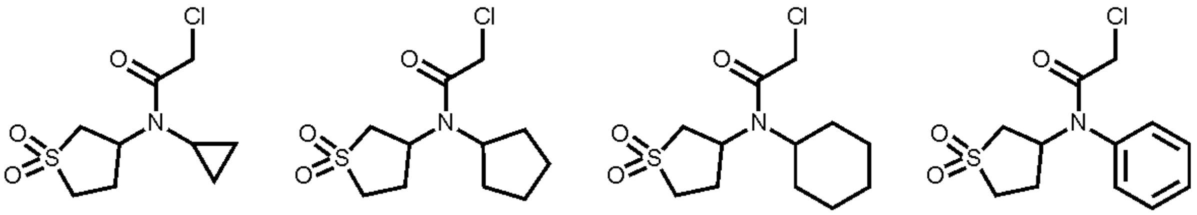

PCM-0102832 PCM-0102313 PCM-0102760 PCM-0102755

**Matched molecular pairs with improved labeling under the same conditions**

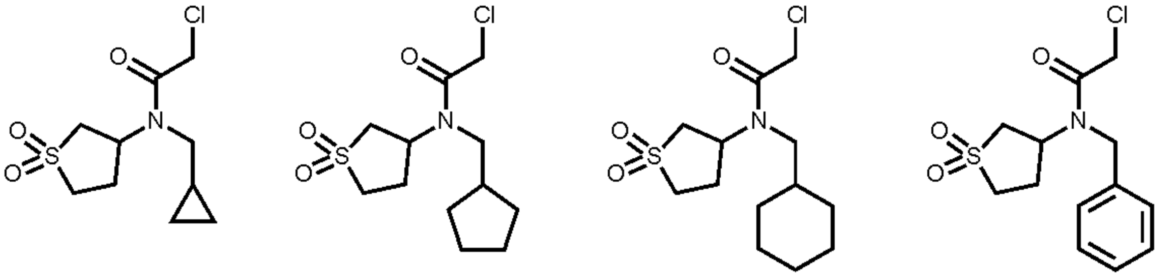

Pin1-3-13
46% labeling

Pin1-3-14
27% labeling

Pin1-2-3
65% labeling

Pin1-437
27% labeling

**Supplementary Figure 4. Matched molecular pair analysis highlights the importance of a methylene linker between the amide and lipophilic moiety.**

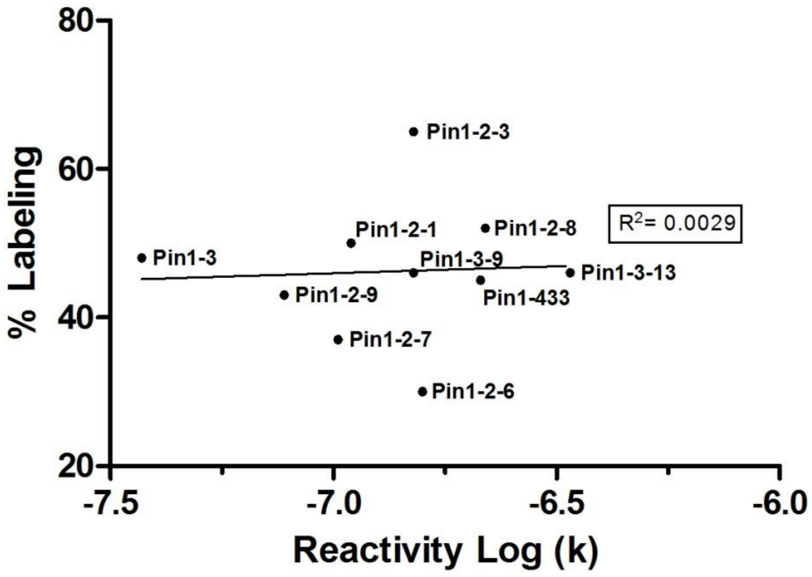


**Supplementary Figure 5. Correlation between labeling percentage and reactivity**
% Labeling and reactivity (log (k)) of the top ten binders shows no correlation ($R^2$ = 0.003).

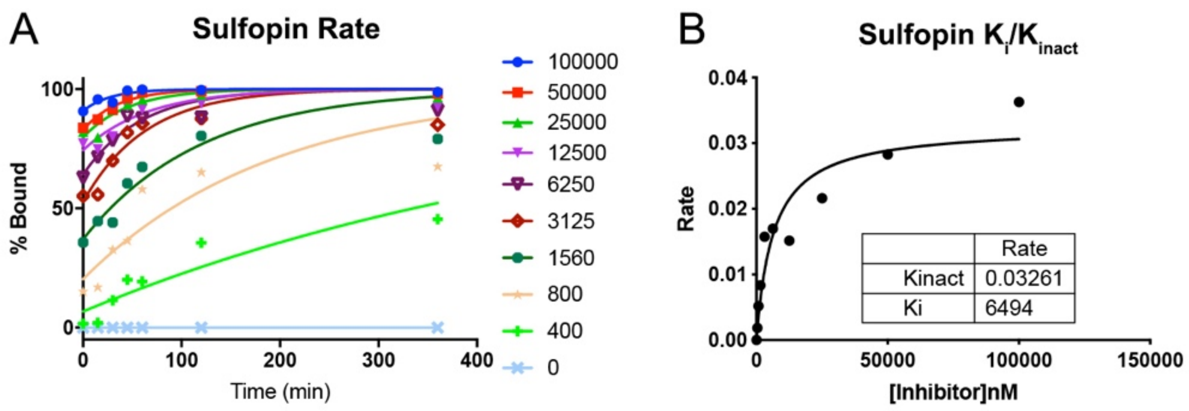


**Supplementary Figure 6. Determination of Sulfopin $K_{inact}/K_i$.** A. Determining labeling rates for various concentrations of Sulfopin. B. Plotting rates as a function of Sulfopin concentration to extrapolate $K_{inact}$ and $K_i$.

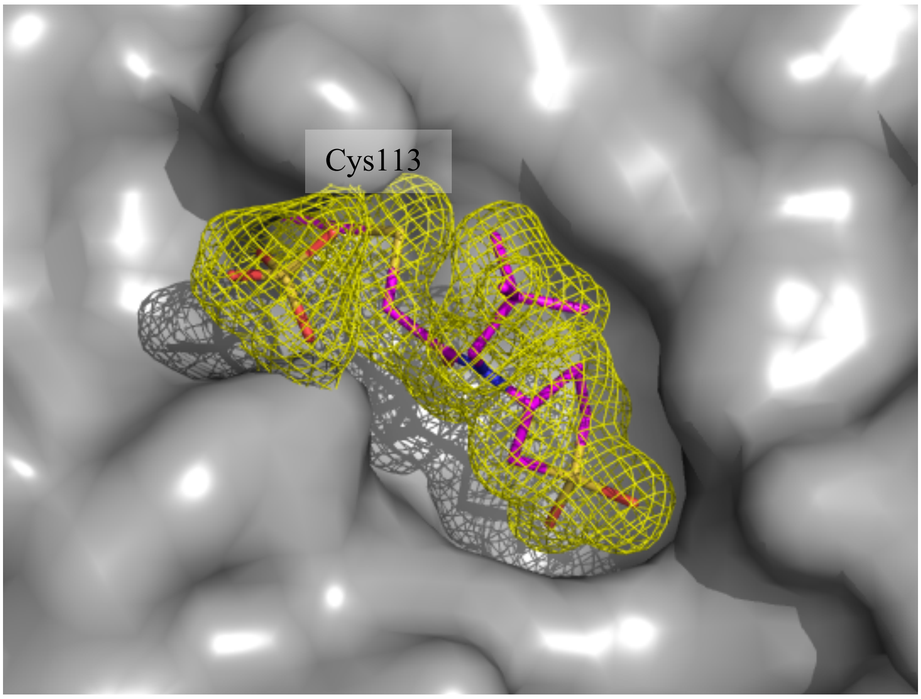


**Supplementary Figure 7**. Continuous electron density between Cys113 and Sulfopin confirms the formation of the covalent bond.

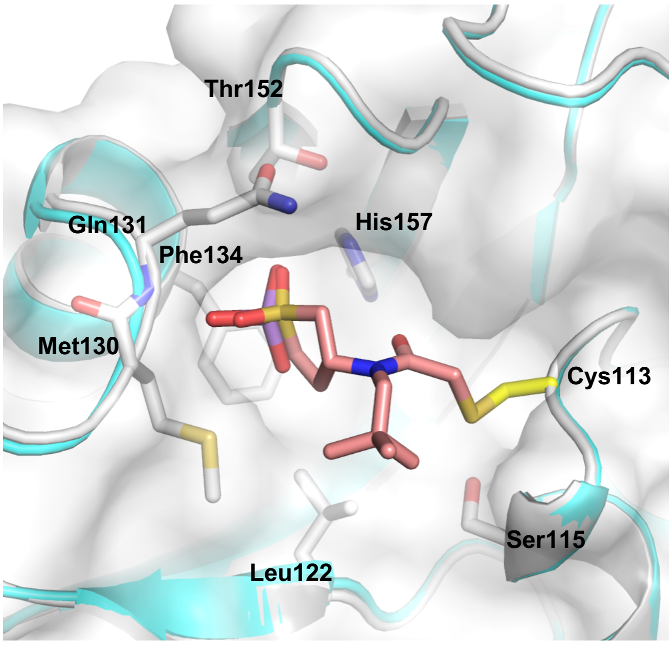


**Supplementary Figure 8. Superposition of Pin1 (white) in complex with Sulfopin (pink; PDB: 6VAJ, 1.4Å) and Pin1 (cyan) in complex with arsenic trioxide (ATO, purple) (PDB: 6DUN, 1.6 Å).**

The sulfolane moiety of Sulfopin and ATO occupy the hydrophobic Pro-binding pocket formed by M130, Q131, F134, Thr152 and H157. The sulfonyl oxygens (red) of Sulfopin and ATO analogously mediate hydrogen bonds with the backbone amide of Q131 and the imidazole NH of H157.

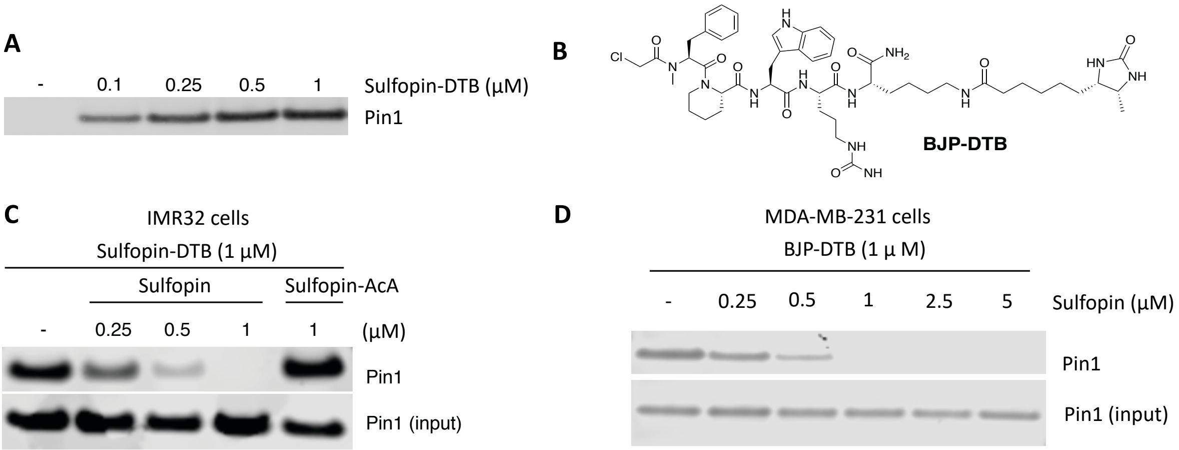


**Supplementary Figure 9. Lysate and cellular engagement of Sulfopin-DTB and Sulfopin**
**A.** Sulfopin-DTB engages Pin1 in PATU-8988T cell lysates.
**B**. BJP-DTB is a desthiobiotin probe based on our recently reported covalent peptide Pin1 inhibitior.
**C.** Sulfopin engages Pin1 in IMR32 cells.
**D.** Sulfopin engages Pin1 in MDA-MB-231 cells.

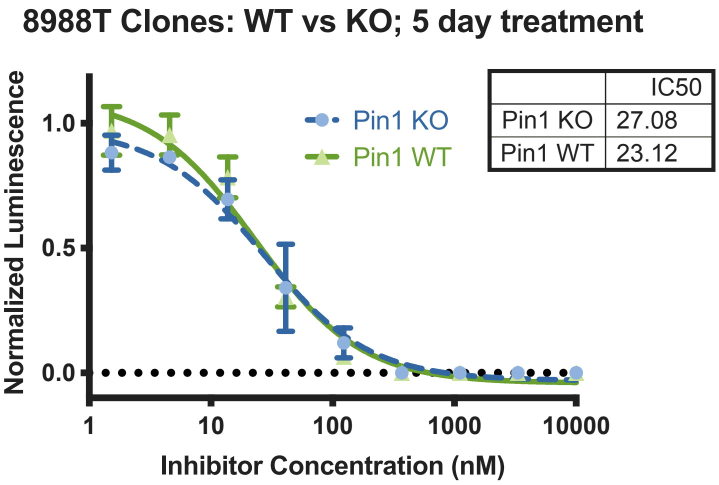


**Supplementary Figure 10. Juglone's viability effects are not Pin1-mediated.** PATU 8988T cells either WT for Pin1 or KO (see Fig. 4A) were treated with Juglone for 5 days. The similar $IC_{50}$ indicates that Juglone's viability effects are Pin1 independent.

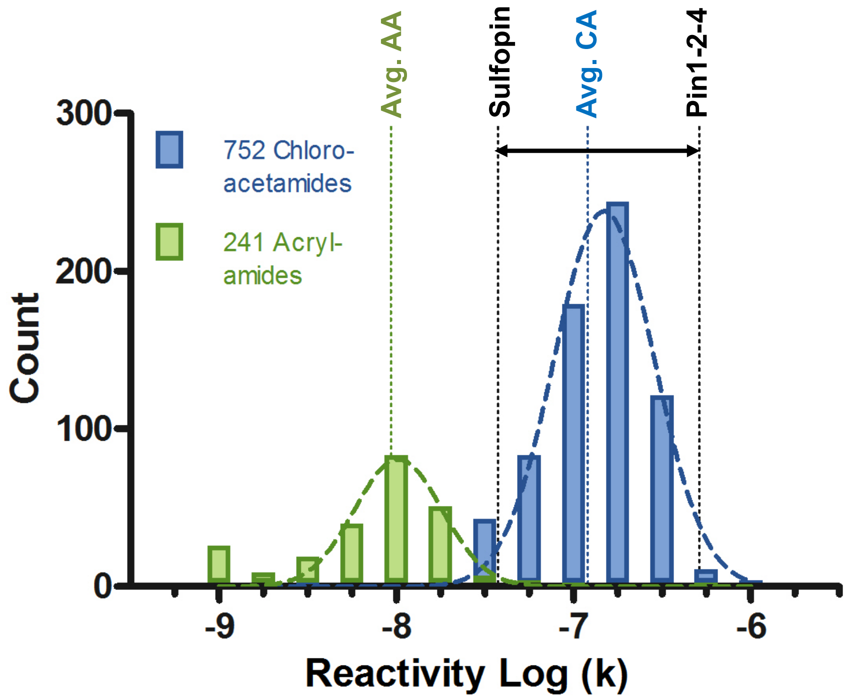


**Supplementary Figure 11. Reactivity distribution of optimized Pin1 covalent inhibitors compared to the electrophile fragment library.** The most reactive Pin1-2-4 is ~14x more reactive than the least reactive of the top analogs - Sulfopin, which shows a reactivity close to an acrylamide.

**Supplementary Tables**

| Compound | Labeling [%] |
|---|---|
| PCM-0102372 | 100 |
| PCM-0102760 | 100 |
| PCM-0102539 | 100 |
| PCM-0102579 | 100 |
| PCM-0102868 | 100 |
| PCM-0102230 | 87 |
| PCM-0102105 | 85 |
| PCM-0102755 | 83 |
| PCM-0102313 | 83 |
| PCM-0102178 | 72 |
| PCM-0102832 | 72 |
| PCM-0103082 | 69 |
| PCM-0102138 | 56 |
| PCM-0102896 | 42 |

**Supplementary Table 1. Labeling of Pin1 by fragments of the electrophilic library that contain 2-sulfolene or sulfolane scaffolds.**
2 µM of Pin1 were incubated with compounds at 200 µM concentration for 24 h at RT. Labeling in % was assigned via intact protein LC/MS. See Supp. Dataset 1 for structures of compounds.

| Compound | Labeling [%][a] | Reactivity k [M^-1*s^-1][b] | Reactivity Log k[c] |
|---|---|---|---|
| Pin1-2-3 | 65 | 1.53E-07 | -6.82 |
| Pin1-2-8 | 52 | 2.19E-07 | -6.66 |
| Pin1-2-1 | 50 | 1.09E-07 | -6.96 |
| Pin1-3 | 48 | 3.73E-08 | -7.43 |
| Pin1-3-13 | 46 | 1.50E-07 | -6.82 |
| Pin1-3-9 | 46 | 3.42E-07 | -6.47 |
| Pin1-433 | 45 | 2.13E-07 | -6.67 |
| Pin1-2-9 | 43 | 7.68E-08 | -7.11 |
| Pin1-2-7 | 37 | 1.02E-07 | -6.99 |
| Pin1-3-7 | 36 | 1.12E-07 | -6.95 |
| Pin1-2-6 | 30 | 1.58E-07 | -6.80 |
| Pin1-053 | 28 | 1.24E-07 | -6.91 |
| Pin1-2-2 | 27 | 8.06E-08 | -7.09 |
| Pin1-3-14 | 27 | 7.03E-08 | -7.15 |
| Pin1-437 | 27 | 1.51E-07 | -6.82 |
| Pin1-128 | 25 | 1.47E-07 | -6.83 |
| Pin1-2-10 | 25 | 1.30E-07 | -6.89 |
| Pin1-2-5 | 24 | 1.31E-07 | -6.88 |
| Pin1-3-8 | 23 | 8.22E-08 | -7.09 |
| Pin1-3-15 | 21 | 7.77E-08 | -7.11 |
| Pin1-2-11 | 19 | 1.15E-07 | -6.94 |
| Pin1-838 | 16 | 1.41E-07 | -6.85 |
| Pin1-028 | 16 | 1.59E-07 | -6.80 |
| Pin1-324 | 12 | 1.59E-07 | -6.80 |
| Pin1-707 | 0 | 1.17E-09 | -8.93 |
| PCM-0102138 | 0 | 1.20E-07 | -6.92 |
| PCM-0102178 | 0 | 1.30E-07 | -6.89 |
| PCM-0102105 | 0 | 1.10E-07 | -6.96 |
| PCM-0102832 | 0 | 6.02E-08 | -7.22 |
| PCM-0102313 | 0 | 1.07E-07 | -6.97 |
| PCM-0102760 | 0 | 1.00E-07 | -7.00 |
| PCM-0102755 | 0 | 1.54E-07 | -6.81 |
| PCM-0102230 | 0 | 8.87E-08 | -7.05 |

**Supplementary Table 2. Characterization of sulfolane analogs and primary screening hits.**
[a] 2 μM Pin1 were incubated with compounds at 2 μM concentration for 1 h at RT. Labeling in % was assigned via intact protein LC/MS. See Supp. Fig. 1 & 3 for structures of compounds.
[b] The second-order rate constant for thiol reactivity as determined by a DTNB assay.
[c] $Log_{10}$ value of the reactivity rate constant.

| Compound | Labeling [%] | Ki [nM][a] | Reactivity k [M^-1*s^-1] | Log k | $EC_{50}$ IMR90 [μM][b] |
|---|---|---|---|---|---|
| Pin1-2-3 | 65 | 17/46 | 1.53E-07 | -6.82 | 7.5 |
| Pin1-2-8 | 52 | 133 | 2.19E-07 | -6.66 | 5.1 |
| Pin1-2-1 | 50 | 7\58 | 1.09E-07 | -6.96 | 2.8 |
| Pin1-3 | 48 | 17/20/56/110 | 3.73E-08 | -7.43 | >25 |
| Pin1-3-13 | 46 | 121 | 1.50E-07 | -6.82 | N.D. |
| Pin1-3-9 | 46 | 411 | 3.42E-07 | -6.47 | N.D. |
| Pin1-433 | 45 | 40/194 | 2.13E-07 | -6.67 | 8.9 |
| Pin1-2-9 | 43 | 83 | 7.68E-08 | -7.11 | 11.3 |
| Pin1-2-7 | 37 | 39 | 1.02E-07 | -6.99 | 6.1 |
| Pin1-2-6 | 30 | 194 | 1.58E-07 | -6.80 | 5.6 |

**Supplementary Table 3. Activity of the top ten labeling compounds.**

[a] Ki values were determined in FP assays after 14 h incubation with compound. Multiple values represent results from multiple independent repetitions of the assay.

[b] $EC_{50}$ values are from a CellTiterGlo viability experiment (48h) with IMR90 cells.

See Supp. Fig. 3 for structures of compounds.

**Supplementary Table 4. Crystallization conditions and data collection and refinement statistics for crystal structures**

| | |
|---|---|
| **RCSB accession code** | 6VAJ |
| **Data collection** [a] | |
| Space group | P $4_3$ $2_1$ 2 |
| Cell dimensions | |
| *a*, *b*, *c* (Å) | 48.96 48.96 137.04 |
| a, b, g (°) | 90.00 90.00 90.00 |
| Resolution (Å) | 39.84 - 1.42 (1.471 - 1.42) [b] |
| $R_{pim}$ | 0.01849 (0.5658) |
| Redundancy | 6.2 (6.3) |
| Completeness (%) | 99.38 (99.72) |
| $I / \sigma I$ | 17.67 (1.54) |
| **Structure solution** | |
| PDB entries used for molecular replacement | 1PIN |
| **Refinement** | |
| No. reflections | 32262 (3163) |
| $R_{work}$ | 0.1923 (0.3278) |
| $R_{free}$ | 0.2144 (0.3227) |
| No. atoms | 1384 |
| Macromolecules | 1229 |
| Ligand/ion | 23 |
| Water | 132 |
| *B*-factors | 31.41 |
| Macromolecules | 30.11 |
| Ligand/ion | 50.67 |
| Water | 40.23 |
| R.m.s. deviations | |
| Bond lengths (Å) | 0.006 |
| Bond angles (°) | 1.19 |
| **Ramachandran** | |
| Preferred | 100.0% |
| Allowed | 0.0% |
| Not Allowed | 0.0% |

[a] A single crystal was used to collect data for each of the structures reported here.

[b] Values in parentheses are for highest-resolution shell.

### Supplementary methods

*Synthetic chemistry methods*

All solvents and reagents used for organic synthesis were purchased from Sigma-Aldrich, Merck, Baker, Acros and used without further purification. Building blocks for synthesis were purchased from Enamine and MolPort chemical suppliers. Purification of precursors was performed using an automated Flash chromatography system (CombiFlash® Systems, Teledyne Isco, USA) with RediSep Rf Normal-phase Flash Columns. Final compounds were purified by semi-preparative HPLC on a Waters Prep 2545 Preparative Chromatography System, with UV/Vis detector 2489, using XBridge® Prep C18 10μm 10x250 mm Column (PN: 186003891, SN:161I3608512502). Reaction progress was monitored using a Waters UPLC-MS system: Acquity UPLC® H class with PDA detector, Acquity UPLC® BEH C18 1.7 μm 2.1x50 mm Column (PN:186002350, SN 02703533825836), Waters SQ detector 2. Spectral analysis by $^{1}H$- and $^{13}C$-NMR was obtained on a Bruker Avance -300 MHz and 400 MHz spectrometer, equipped with a QNP probe. Chemical shifts ($\delta_H$ & $\delta_C$) are quoted in ppm to the nearest 0.1 ppm, and referenced to trimethylsilane (TMS). Coupling constants (J) are reported in Hertz (Hz) to the nearest 0.1 Hz. High resolution electron-spray mass spectrometry (HRMS-ESI) was performed on a Xevo G2-XS QTOF Mass Spectrometer (Waters Corporation, USA) and reported values are within the error limit of ± 2.2 ppm mass units.

*General procedure for chloroacetamide preparation*

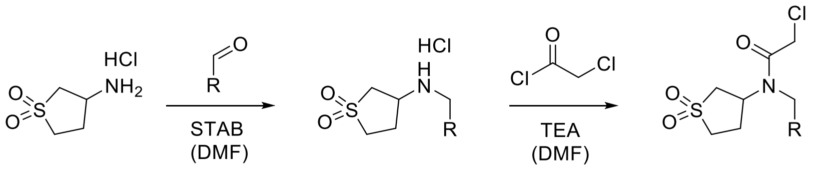


3-aminosulfolane hydrochloride (1 eq.) was added to a solution of triethylamine (TEA) (0.9 eq.) in dry dimethylformamide (DMF) and stirred for 1 h at room temperature (RT). Afterwards, aldehyde (1.1 eq.) and acetic acid (0.2 eq.) were added to the reaction mixture and stirred at RT for 1 h. Then sodium triacetoxyborohydride (STAB) (2.1 eq.) was added at once to the mixture and stirred overnight at RT. After evaporation of the solvent, the residue was dissolved with sat. aq. $NaHCO_3$ and the aq. solution was extracted with ethyl acetate (EA) (2x). The organic layers were combined, dried over $Na_2SO_4$ and filtered. Evaporation of the solvent yielded the secondary amine as hydrochloride, which was used without purification in the next step. Secondary amine hydrochloride (1 eq.) was dissolved in dry DMF and cooled to 0 °C. Subsequently, 2-chloroacetyl chloride (1.2 eq.) and TEA (1.2 eq.) were added dropwise at 0 °C and stirred for 30 min. Afterwards the reaction mixture was allowed to reach RT and stirred for 1 h. The reaction was quenched at 0 °C by the addition of water and filtered. Purification by RP-HPLC (linear gradient 5 → 95% ACN/$H_2O$ + 0.1% TFA in 30 min) and lyophilization yielded the corresponding chloroacetamide.

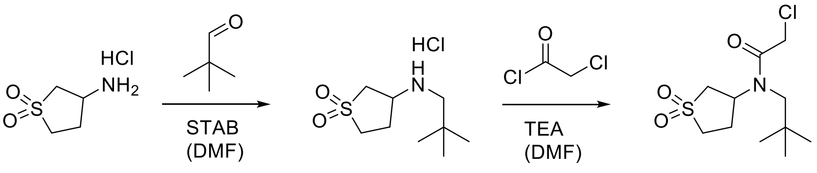

1 Pin1-3

**2-chloro-*N*-(sulfolan-3-yl)-*N*-neopentylacetamide (Sulfopin / Pin1-3)**

Following the general procedure of chloroacetamide preparation, 3-aminosulfolane hydrochloride (100 mg, 0.583 mmol, 1 eq.) was added to a solution of triethylamine (TEA) (73 µL, 0.524 mmol, 0.9 eq.) in dry dimethylformamide (DMF) (1.4 mL) and stirred for 1 h at room temperature (RT). Afterwards, pivaldehyde (70 µL, 0.641 mmol, 1.1 eq.) and one drop of acetic acid (0.2 eq.) were added to the reaction mixture and stirred at RT for 1 h. Then sodium triacetoxyborohydride (STAB) (259 mg, 1.223 mmol, 2.1 eq.) was added at once to the mixture and stirred overnight at RT. After evaporation of the solvent, the residue was dissolved with sat. aq. $NaHCO_3$ (0.5 mL) and the aq. solution was extracted with ethyl acetate (EA) (2x 1 mL). The organic layers were combined, dried over $Na_2SO_4$ and filtered. Evaporation of the solvent yielded the secondary amine **1** as white solid (86.2 mg, 0.42 mmol, 72% (crude product)), which was used without purification in the next step.

Secondary amine **1** hydrochloride (100 mg, 0.487 mmol, 1 eq.) was dissolved in dry DMF (1 mL) and cooled to 0 °C. Subsequently, 2-chloroacetyl chloride (47 µL, 0.584 mmol, 1.2 eq.) and TEA (81 µL, 0.584 mmol, 1.2 eq.) were added dropwise at 0 °C and stirred for 30 min. Afterwards the reaction mixture was allowed to reach RT and stirred for 2 h. The reaction was quenched at 0 °C by the addition of water (2 mL). Purification by RP-HPLC ($t_R$ = 16 min, linear gradient 5 → 95% ACN/$H_2O$ + 0.1% TFA in 30 min) and lyophilization yielded chloroacetamide Pin1-3 (59.8 mg, 0.212 mmol, 44%) as white powder.

**$^1$H NMR** (500 MHz, $CDCl_3$): δ = 4.11 (d, *J*=5.5 Hz, 2 H), 3.91 - 4.00 (m, 1 H), 3.65 - 3.79 (m, 2 H), 3.25 - 3.33 (m, 1 H), 3.11 - 3.20 (m, 2 H), 3.00 - 3.09 (m, 1 H), 2.46 - 2.60 (m, 2 H), 1.03 (s, 9 H) ppm.
**$^{13}$C NMR** (126 MHz, $CDCl_3$): δ = 168.0, 62.4, 57.6, 50.3, 49.0, 42.1, 33.6, 28.0, 26.6 ppm.
**HRMS** (ESI): *m/z* calcd. for $C_{11}H_{20}ClNO_3SNa^+$ $[M+Na]^+$: 304.0750; found 304.0746.

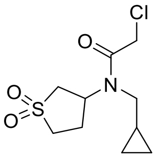

Pin1-3-13

**2-chloro-*N*-(cyclopropylmethyl)-*N*-(sulfolan-3-yl)acetamide (Pin1-3-13)**

Following the general procedure of chloroacetamide preparation, 3-aminosulfolane hydrochloride (100 mg, 0.583 mmol, 1 eq.), TEA (73 µL, 0.524 mmol, 0.9 eq.) in dry DMF (1.4 mL) were stirred for 1 h at RT. Cyclopropanecarboxaldehyde (48 µL, 0.641 mmol, 1.1 eq.), one drop of acetic acid and STAB (259 mg, 1.223 mmol, 2.1 eq.) were added and stirred overnight at RT. After workup and evaporation, the secondary amine as hydrochloride (63.1 mg, 0.280 mmol, 48% (crude product)) was used without purification in the next step. 2-chloroacetyl chloride (27 µL, 0.335 mmol, 1.2 eq.) and TEA (47 µL, 0.335 mmol, 1.2 eq.) were added dropwise to cooled (0 °C) secondary amine hydrochloride (63.1 mg, 0.280 mmol, 1 eq.) in dry DMF (0.6 mL) and stirred for 30 min. Quenching with water (2 mL), filtering, purification by RP-HPLC ($t_R$ = 13.5

min, linear gradient 5 → 95% ACN/$H_2O$ + 0.1% TFA in 30 min) and lyophilization yielded chloroacetamide Pin1-3-13 (24.9 mg, 0.094 mmol, 34%) as white powder.

$^{1}$**H NMR** (500MHz, $CDCl_3$): δ = 4.27 - 4.41 (m, 1 H), 4.13 (s, 2 H), 3.52 - 3.73 (m, 2 H), 3.17 - 3.38 (m, 3 H), 3.02 - 3.14 (m, 1 H), 2.45 - 2.61 (m, 2 H), 0.91 - 1.04 (m, 1 H), 0.73 (d, *J*=6.6 Hz, 2 H), 0.36 (d, *J*=4.4 Hz, 2 H) ppm.
$^{13}$**C NMR** (126MHz, $CDCl_3$): δ = 166.7, 54.4, 54.1, 50.7, 50.2, 41.9, 26.4, 11.1, 4.3 ppm.
**HRMS** (ESI): *m/z* calcd. for $C_{10}H_{16}ClNO_3SNa^+$ $[M+Na]^+$: 288.0437; found 288.0435.

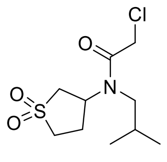

Pin1-3-15

**2-chloro-*N*-(sulfolan-3-yl)-*N*-isobutylacetamide (Pin1-3-15)**

Following the general procedure of chloroacetamide preparation, 3-aminosulfolane hydrochloride (90 mg, 0.524 mmol, 1 eq.), TEA (66 µL, 0.474 mmol, 0.9 eq.) in dry DMF (1.3 mL) were stirred for 1 h at RT. Isobutyraldehyde (57 µL, 0.629 mmol, 1.2 eq.), one drop of acetic acid (0.2 eq.) and STAB (233 mg, 1.101 mmol, 2.1 eq.) were added and stirred overnight at RT. After workup and evaporation, the secondary amine as hydrochloride (78.18 mg, 0.343 mmol, 66% (crude product)) was used without purification in the next step. 2-chloroacetyl chloride (33 µL, 0.412 mmol, 1.2 eq.) and TEA (57 µL, 0.412 mmol, 1.2 eq.) were added dropwise to cooled (0 °C) secondary amine hydrochloride (78.18 mg, 0.487 mmol, 1 eq.) in dry DMF (0.7 mL) and stirred for 30 min. Quenching with water (2 mL), filtering, purification by RP-HPLC ($t_R$ = 14 min, linear gradient 5 → 95% ACN/$H_2O$ + 0.1% TFA in 30 min) and lyophilization yielded chloroacetamide Pin1-3-15 (29.22 mg, 0.412 mmol, 32%) as white powder.

$^{1}$**H NMR** (500 MHz, $CDCl_3$): δ = 4.10 (s, 2 H), 3.99 - 4.08 (m, 1 H), 3.56 - 3.77 (m, 2 H), 3.11 - 3.22 (m, 3 H), 3.01 - 3.10 (m, 1 H), 2.44 - 2.60 (m, 2 H), 1.86 - 1.98 (m, 1 H), 0.99 (t, *J*=6.6 Hz, 6 H) ppm.
$^{13}$**C NMR** (126 MHz, $CDCl_3$): δ = 167.0, 58.0, 55.2, 50.5, 49.7, 42.0, 28.4, 26.2, 19.9, 19.7 ppm
**HRMS** (ESI): *m/z* calcd. for $C_{10}H_{18}ClNO_3SNa^+$ $[M+Na]^+$: 209.0594; found 290.0590.

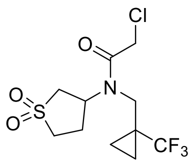

Pin1-3-9

**2-chloro-*N*-(sulfolan-3-yl)-*N*-((1-(trifluoromethyl)cyclopropyl)methyl)acetamide (Pin1-3-9)**

Following the general procedure of chloroacetamide preparation, 3-aminosulfolane hydrochloride (55 mg, 0.320 mmol, 1 eq.), TEA (40 µL, 0.288 mmol, 0.9 eq.) in dry DMF (0.8 mL) were stirred for 1 h at RT. 1-(Trifluoromethyl)cyclopropanecarbaldehyde (48.7 mg, 0.352 mmol, 1.1 eq.), one drop of acetic acid and STAB (143 mg, 0.673 mmol, 2.1 eq.) were added and stirred overnight at RT. After workup and evaporation, the secondary amine as hydrochloride (72 mg, 0.245 mmol, 76% (crude product)) was used without purification in the next step. 2-chloroacetyl chloride (22 µL, 0.270 mmol, 1.1 eq.) and TEA (41 µL, 0.294 mmol, 1.2 eq.) were added dropwise to cooled (0 °C) secondary amine hydrochloride (72 mg, 0.245 mmol, 1 eq.) in dry DMF (0.6 mL) and stirred for 30 min. Quenching with water (2 mL), filtration, purification by RP-HPLC ($t_R$ = 16.5 min, linear gradient 5 → 95% ACN/$H_2O$ + 0.1% TFA in 30 min) and lyophilization yielded chloroacetamide Pin1-3-9 (18.8 mg, 0.056 mmol, 23%) as white powder.

**$^1$H NMR** (500MHz, $CDCl_3$): δ = 4.11 - 4.18 (m, 2 H), 3.98 - 4.10 (m, 1 H), 3.57 - 3.73 (m, 4 H), 3.10 - 3.22 (m, 1 H), 3.02 - 3.09 (m, 1 H), 2.47 - 2.56 (m, 2 H), 1.18 - 1.28 (m, 2 H), 0.83 - 0.91 (m, 2 H) ppm.
**$^{13}$C NMR** (126 MHz, $CDCl_3$): δ = 167.2, 127.4, 125.2, 55.3, 51.9, 50.4, 49.3, 41.5, 26.0, 25.4, 8.3 ppm.
**HRMS** (ESI): *m/z* calcd. for $C_{11}H_{15}ClNO_3SNa^+$ $[M+Na]^+$: 356.0311; found 356.0306.

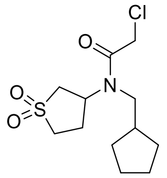


Pin1-3-14

**2-chloro-*N*-(cyclopentylmethyl)-*N*-(sulfolan-3-yl)acetamide (Pin1-3-14)**

Following the general procedure of chloroacetamide preparation, 3-aminosulfolane hydrochloride (100 mg, 0.583 mmol, 1 eq.), TEA (73 µL, 0.524 mmol, 0.9 eq.) in dry DMF (1.4 mL) were stirred for 1 h at RT. Cyclopentanecarboxaldehyde (68 µL, 0.641 mmol, 1.1 eq.), one drop of acetic acid and STAB (259 mg, 1.223 mmol, 2.1 eq.) were added and stirred overnight at RT. After workup and evaporation, the secondary amine (95.68 mg, 0.377 mmol, 65% (crude product)) was used without purification in the next step. 2-chloroacetyl chloride (36 µL, 0.452 mmol, 1.2 eq.) and TEA (63 µL, 0.452 mmol, 1.2 eq.) were added dropwise to cooled (0 °C) secondary amine hydrochloride (95.68 mg, 0.377 mmol, 1 eq.) in dry DMF (1 mL) and stirred for 30 min. Quenching with water (2 mL), filtration, purification by RP-HPLC ($t_R$ = 17.5 min, linear gradient 5 → 95% ACN/$H_2O$ + 0.1% TFA in 30 min) and lyophilization yielded Pin1-3-14 (23.4 mg, 0.08 mmol, 21%) as white powder.

**$^1$H NMR** (500 MHz, $CDCl_3$): δ = 4.11 (s, 3 H), 3.57 - 3.76 (m, 2 H), 3.26 - 3.39 (m, 2 H), 3.02 - 3.20 (m, 2 H), 2.47 - 2.58 (m, 2 H), 2.11 - 2.21 (m, 1 H), 1.77 - 1.88 (m, 2 H), 1.60 - 1.76 (m, 4 H), 1.17 - 1.29 (m, 2 H) ppm.
**$^{13}$C NMR** (125 MHz, $CDCl_3$): δ = 166.8, 55.3, 55.1, 50.5, 49.7, 42.0, 40.2, 30.4, 30.4, 26.3, 24.9, 24.9 ppm.
**HRMS** (ESI): *m/z* calcd. for $C_{12}H_{20}ClNO_3SNa^+$ $[M+Na]^+$: 316.0750; found 316.0743.

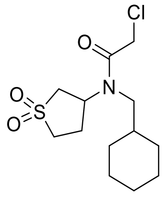


Pin1-2-3

**2-chloro-*N*-(cyclohexylmethyl)-*N*-(sulfolan-3-yl)acetamide (Pin1-2-3)**

Following the general procedure of chloroacetamide preparation, 3-aminosulfolane hydrochloride (75 mg, 0.437 mmol, 1 eq.) in dry DMF (1.1 mL) were stirred for 1 h at RT. Cyclohexanecarboxaldehyde (58 µL, 0.481 mmol, 1.1 eq.) and STAB (139 mg, 0.655 mmol, 1.5

eq.) were added and stirred overnight at RT. After workup and evaporation, the secondary amine as hydrochloride (72.11 mg, 0.269 mmol, 62% (crude product)) was used without purification in the next step. 2-chloroacetyl chloride (25 µL, 0.323 mmol, 1.2 eq.) and TEA (45 µL, 0.323 mmol, 1.2 eq.) were added dropwise to cooled (0 °C) secondary amine hydrochloride (72 mg, 0.269 mmol, 1 eq.) in dry DMF (0.6 mL) and stirred for 30 min. Quenching with water (2 mL), filtration, purification by RP-HPLC ($t_R$ = 18.5 min, linear gradient 5 → 95% ACN/$H_2O$ + 0.1% TFA in 30 min) and lyophilization yielded Pin1-2-3 (9.1 mg, 0.030 mmol, 11%) as white powder.

**$^1$H NMR** (500 MHz, $CDCl_3$): δ = 4.09 (s, 2 H), 4.00 - 4.07 (m, 1 H), 3.56 - 3.77 (m, 2 H), 3.01 - 3.27 (m, 4 H), 2.45 - 2.59 (m, 2 H), 1.66 - 1.88 (m, 5 H), 1.51 - 1.63 (m, 1 H), 1.13 - 1.33 (m, 3 H), 0.89 - 1.03 (m, 2 H) ppm.
**$^{13}$C NMR** (125 MHz, $CDCl_3$): δ = 167.0, 57.1, 55.3, 50.5, 49.7, 42.0, 38.0, 30.9, 30.8, 26.2, 26.1, 25.8 ppm.
**HRMS** (ESI): *m/z* calcd. for $C_{13}H_{22}ClNO_3SNa^+$ $[M+Na]^+$: 330.0907; found 330.0901.

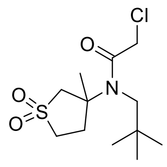


**Pin1-18**

**2-chloro-*N*-(3-methyl-sulfolan-3-yl)-*N*-neopentylacetamide (Pin1-18)**

Following the general procedure of chloroacetamide preparation, 3-amino-3-methylsulfolane hydrochloride (200 mg, 1.077 mmol, 1 eq.), TEA (135 µL, 0.969 mmol, 0.9 eq.) in dry DMF (2.7 mL) were stirred for 1 h at RT. Pivaldehyde (129 µL, 1.185 mmol, 1.1 eq.), two drops of acetic acid and STAB (479 mg, 2.262 mmol, 2.1 eq.) were added and stirred overnight at RT. After workup and evaporation, the secondary amine (267.6 mg, 1.046 mmol, 97% (crude product)) was used without purification in the next step. 2-chloroacetyl chloride (17.98 µL, 0.235 mmol, 1.2 eq.) and TEA (32.7 µL, 0.235 mmol, 1.2 eq.) were added dropwise to cooled (0 °C) secondary amine hydrochloride (50 mg, 0.195 mmol, 1 eq.) in dry DMF (0.4 mL) and stirred for 30 min. Quenching with water (2 mL) and purification by RP-HPLC ($t_R$ = 17 min, linear gradient 5 → 95% ACN/$H_2O$ + 0.1% TFA in 30 min) yielded Pin1-18 (11.94 mg, 0.04 mmol, 21%) as white powder.

**$^1$H NMR** (500 MHz, $CDCl_3$): δ = 3.96 - 4.44 (m, 2 H), 3.68 (d, *J*=13.2 Hz, 1 H), 3.51 - 3.65 (m, 1 H), 3.41 (s, 2 H), 3.12 - 3.37 (m, 2 H), 2.13 - 2.54 (m, 2 H), 1.77 (s, 3 H), 1.04 (s, 9 H) ppm.
**$^{13}$C NMR** (126 MHz, $CDCl_3$): δ = 170.0, 60.9, 55.0, 51.0, 43.5, 33.7, 29.1, 27.1 ppm.
**HRMS** (ESI): *m/z* calcd. for $C_{12}H_{22}ClNO_3SNa^+$ $[M+H^+]$: 318.0907; found 318.0914.

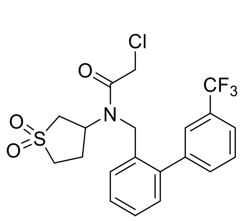


**Pin1-3-8**

**2-chloro-*N*-(sulfolan-3-yl)-*N*-((3'-(trifluoromethyl)-[1,1'-biphenyl]-2-yl)methyl)acetamide (Pin1-3-8)**

3-aminosulfolane hydrochloride (100 mg, 0.583 mmol, 1 eq.) was added to a solution of TEA (73.1 µL, 0.524 mmol, 0.9 eq.) in MeOH (2 mL) and stirred for 1 h at room temperature (RT). Afterwards, 3'-(trifluoromethyl)-1,1'-biphenyl-2-carboxaldehyde (182 µL, 0.728 mmol, 1.25 eq.) and acetic acid (6.67 µL, 0.117 mmol, 0.2 eq.) were added to the reaction mixture and stirred at RT for 1 h. Then sodium cyanoborohydride (92 mg, 1.456 mmol, 2.5 eq.) was added at once to the mixture and stirred overnight at RT. After evaporation of the solvent, the residue was dissolved with sat. aq. $NaHCO_3$ (0.5 mL) and the aq. solution was extracted with ethyl acetate (EA) (2x 1 mL). The organic layers were combined, dried over $Na_2SO_4$ and filtered. Evaporation of the solvent yielded the secondary amine **1** as white solid (231.8 mg, 0.571 mmol, 98% (crude product)), which was used without purification in the next step. The secondary amine hydrochloride (100 mg, 0.271 mmol, 1 eq.) was dissolved in dry DMF (0.5 mL) and cooled to 0 °C. Subsequently, 2-chloroacetyl chloride (26 µL, 0.325 mmol, 1.2 eq.) and TEA (56.6 µL, 0.406 mmol, 1.5 eq.) were added dropwise at 0 °C and stirred for 30 min. Afterwards the reaction mixture was allowed to reach RT and stirred for 2 h. The reaction was quenched at 0 °C by the addition of water (2 mL). Purification by RP-HPLC ($t_R$ = 22 min, linear gradient 5 → 95% ACN/$H_2O$ + 0.1% TFA in 30 min) and lyophilization yielded chloroacetamide Pin1-3-8 (40.69 mg, 0.091 mmol, 34%) as white powder.

**$^1$H NMR** (500 MHz, CDCl3): δ = 7.73 (d, J=7.7 Hz, 1 H), 7.65 (t, J=7.7 Hz, 1 H), 7.56 - 7.61 (m, 1 H), 7.43 - 7.56 (m, 3 H), 7.21 - 7.39 (m, 2 H), 5.06 (br. s., 1 H), 4.69 (br. s., 1 H), 4.50 - 4.61 (m, 2 H), 4.42 (br. s., 1 H), 4.20 (s, 1 H), 3.23 - 3.45 (m, 2 H), 3.06 - 3.18 (m, 1 H), 2.95 - 3.05 (m, 1 H), 2.25 - 2.43 (m, 2 H) ppm.

**$^{13}$C NMR** (126 MHz, $CDCl_3$): δ = 167.2, 140.3, 132.2, 131.0, 129.5, 129.2, 129.0, 128.6, 125.7, 124.9, 62.7, 53.0, 50.6, 40.4, 26.3 ppm.

**HRMS** (ESI): *m/z* calcd. for $C_{20}H_{19}ClF_3NO_3SNa^+$ $[M+Na]^+$: 468.0624; found 468.0616.

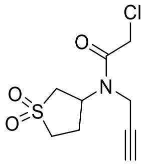


Pin1-4

**2-chloro-*N*-(sulfolan-3-yl)-*N*-(prop-2-yn-1-yl)acetamide (Pin1-4)**

3-(prop-2-yn-1-ylamino)sulfolane hydrochloride (100 mg, 0.477 mmol, 1 eq.) was dissolved in dry DMF (1 mL) and cooled to 0 °C. Subsequently, 2-chloroacetyl chloride (44 µL, 0.572 mmol, 1.2 eq.) and TEA (120 µL, 0.858 mmol, 1.8 eq.) were added dropwise at 0 °C and stirred for 30 min. Afterwards the reaction mixture was allowed to reach RT and stirred for 30 min. The reaction was quenched at 0 °C by the addition of water (2 mL). Subsequent filtration, purification by RP-HPLC ($t_R$ = 11 min, linear gradient 5 → 95% ACN/$H_2O$ + 0.1% TFA in 30 min) and lyophilization yielded chloroacetamide Pin1-4 (52 mg, 0.209 mmol, 44%) as white powder.

**$^1$H NMR** (500 MHz, $CDCl_3$): δ = 4.90 - 5.03 (m, 1 H), 4.16 - 4.25 (m, 4 H), 3.31 - 3.51 (m, 3 H), 3.02 - 3.20 (m, 1 H), 2.39 - 2.89 (m, 3 H) ppm.
**$^{13}$C NMR** (126 MHz, $CDCl_3$): δ = 167.0, 74.6, 52.1, 50.9, 41.5, 35.7, 26.3 ppm.
**HRMS** (ESI): *m/z* calcd. for $C_9H_{12}ClNO_3SNa^+$ $[M+Na]^+$: 272.0124; found 272.0119.

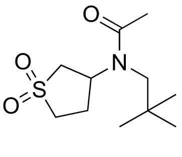

**Sulfopin-AcA**

**_N_-(sulfolan-3-yl)-_N_-neopentylacetamide (Sulfopin-AcA)**

3-(neopentylamino)sulfolane hydrochloride (75 mg, 0.310 mmol, 1 eq.) was dissolved in dry DMF (1 mL) and acetic anhydride (64 µL, 0.682 mmol, 2.2 eq.) and TEA (30 µL, 0.217 mmol, 0.7 eq.) were added dropwise. The reaction was stirred at RT overnight. The reaction was quenched by the addition of water (2 mL). Filtration, purification by RP-HPLC ($t_R$ = 8 min, linear gradient 5 → 95% ACN/$H_2O$ + 0.1% TFA in 30 min) and lyophilization yielded chloroacetamide Sulfopin-AcA (8.75 mg, 0.035 mmol, 34%) as white powder.

**$^1$H NMR** (500 MHz, $CDCl_3$): δ = 3.87 - 3.97 (m, 1 H), 3.75 - 3.84 (m, 1 H), 3.63 - 3.74 (m, 1 H), 3.28 (d, *J*=15.4 Hz, 1 H), 3.07 - 3.15 (m, 2 H), 2.99 - 3.06 (m, 1 H), 2.47 - 2.59 (m, 2 H), 2.12 - 2.18 (m, 3 H), 0.98 - 1.08 (m, 9 H) ppm.
**$^{13}$C NMR** (126 MHz, $CDCl_3$): δ = 172.9, 63.1, 57.0, 50.4, 49.4, 33.8, 28.1, 26.9, 23.2 ppm.
**HRMS** (ESI): *m/z* calcd. for $C_{11}H_{21}NO_3SNa^+$ $[M+Na]^+$: 270.1140; found 270.1140.

*General scheme of the DTB-Probe synthesis*

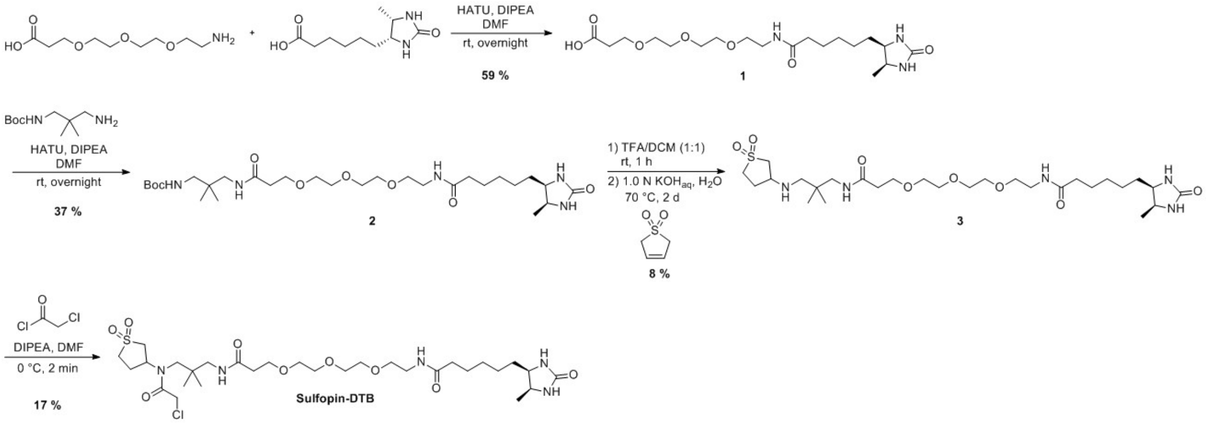


*Experimental Details for Individual Compound Synthesis*

**19-((4R,5S)-5-methyl-2-oxoimidazolidin-4-yl)-14-oxo-4,7,10-trioxa-13-azanonadecanoic acid**

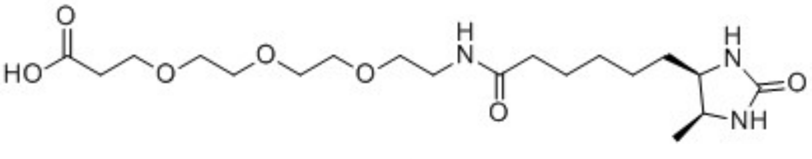

**1**

6-((4R,5S)-5-methyl-2-oxoimidazolidin-4-yl)hexanoic acid (96.8 mg, 0.452 mmol), HATU (171.9 mg, 0.452 mmol) and DIPEA (240 µL, 1.36 mmol) were dissolved in DMF (1.0 mL) and stirred at r.t. for 15 min. The resulting reaction mixture was added dropwise to a solution of 3-(2-(2-(2-aminoethoxy)ethoxy)ethoxy)propanoic acid (100.0 mg, 0.452 mmol) in DMF (1.0 mL) until all the starting material was consumed. The crude was purified by HPLC to yield the product as a yellow oil (142 mg, 0.340 mmol, 59%).

**MS** (ESI): *m/z* calcd. for C19H36N3O7$^+$ $[M+H]^+$: 418.25; found 418.4.

**tert-butyl(2,2-dimethyl-23-((4R,5S)-5-methyl-2-oxoimidazolidin-4-yl)-5,18-dioxo-8,11,14-trioxa-4,17-diazatricosyl)carbamate**

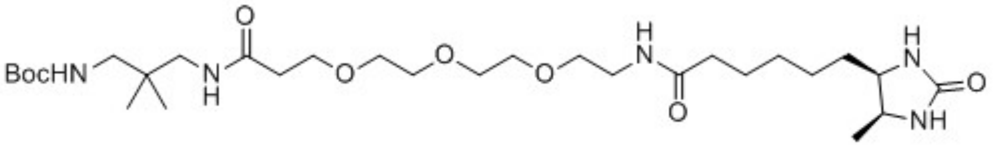

**2**

Intermediate 1 (130.0 mg, 0.311 mmol), tert-butyl (3-amino-2,2-dimethylpropyl)carbamate (75.5 mg, 0.373 mmol), HOBt (75.7 mg, 0.560 mmol) and DIPEA (163 µL, 0.933 mmol) were dissolved in DMF (1.0 mL). EDC·HCl (107.4 mg, 0.560 mmol) was added to the reaction mixture, which was stirred at r.t. for 16 h. The crude was purified by HPLC to yield the product as a dark yellow oil (82 mg, 0.136 mmol, 37%).

**MS** (ESI) *m/z* calcd. for $C_{29}H_{56}N_5O_8^+$ $[M+H]^+$: 602.41; found 602.6.

**N-(16-((1,1-dioxidotetrahydrothiophen-3-yl)amino)-15,15-dimethyl-12-oxo-3,6,9-trioxa-13-azahexadecyl)-6-((4R,5S)-5-methyl-2-oxoimidazolidin-4-yl)hexanamide**

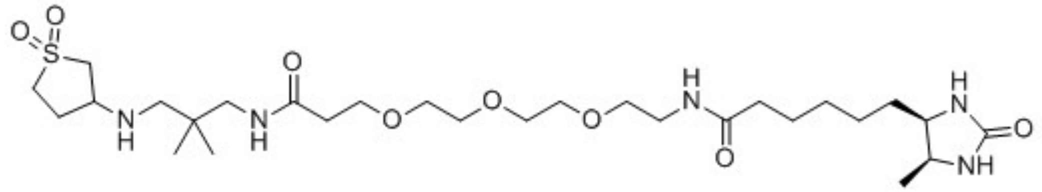

**3**

Intermediate 2 (52.5 mg, 0.0872 mmol) and TFA (1.0 mL) were stirred at r.t. in DCM (1.0 mL) for 1 h. The solvent was removed *in vacuo* and the residue was dissolved in $H_2O$ (0.57 mL). 2,5-dihydrothiophene 1,1-dioxide (17.5 mg, 0.148 mmol) and 1.0 n aq. KOH (500 µL) were added to the solution and stirred at 70 °C for 2 d. The reaction mixture was concentrated *in vacuo* and the residue was purified by HPLC to yield the product as a yellow solid (6 mg, 0.010 mmol, 8%).

**MS** (ESI) *m/z* calcd. for $C_{28}H_{54}N_5O_8S^+$ $[M+H]^+$: 620.37; found 620.5.

**N-(19-chloro-17-(1,1-dioxidotetrahydrothiophen-3-yl)-15,15-dimethyl-12,18-dioxo-3,6,9-trioxa-13,17-diazanonadecyl)-6-((4R,5S)-5-methyl-2-oxoimidazolidin-4-yl)hexanamide**

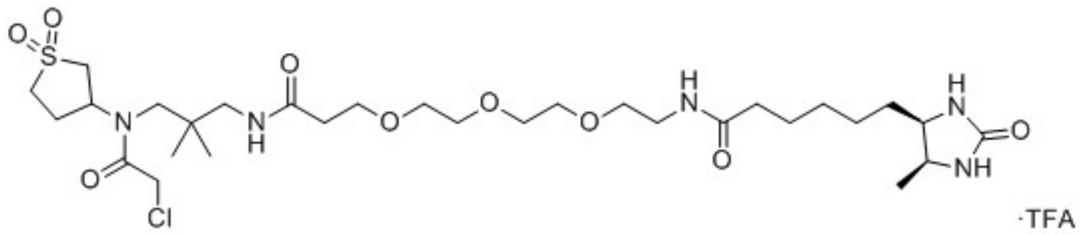


**Sulfopin-DTB**

Intermediate 3 (6 mg, 0.010 mmol) and DIPEA (16 µL, 0.092 mmol) were dissolved in DMF (1.0 mL). The solution was cooled to 0 °C. 2-chloroacetyl chloride (18 µL) in DMF (300 µL) was added dropwise to the reaction mixture until the starting material was consumed. The crude was purified by HPLC to yield the title compound Pin1-3-DTB as a light-yellow solid (1.3 mg, 0.002 mmol, 17%).

**$^1$H NMR** (500 MHz, DMSO-*d*$_6$) δ = 7.81 (s, 1H), 6.28 (s, 1H), 6.11 (s, 1H), 4.33 (s, 1H), 4.02 – 3.92 (m, 1H), 3.61 (t, *J* = 6.3 Hz, 5H), 3.49 (s, 11H), 3.33 (s, 3H), 3.23 (s, 2H), 3.18 (d, *J* = 5.8 Hz, 3H), 3.10 (dt, *J* = 13.2, 6.8 Hz, 3H), 2.99 (s, 2H), 2.38 (s, 3H), 2.09 – 2.02 (m, 3H), 1.46 (d, *J* = 7.3 Hz, 2H), 1.28 (d, *J* = 47.9 Hz, 3H), 0.95 (d, *J* = 6.4 Hz, 3H), 0.88 (s, 6H) ppm.

**MS** (ESI) *m/z* calcd. for $C_{30}H_{55}ClN_5O_9S^+$ $[M+H]^+$: 696.34; found 696.6.
